# *Streptococcus pneumoniae*, *S. mitis*, and *S. oralis* produce a phosphatidylglycerol-dependent, *ltaS*-independent glycerophosphate-linked glycolipid

**DOI:** 10.1101/2020.10.27.355933

**Authors:** Yahan Wei, Luke R. Joyce, Ashley M. Wall, Ziqiang Guan, Kelli L. Palmer

**Affiliations:** Department of Biological Sciences, The University of Texas at Dallas, Richardson, Texas, USA; Department of Biochemistry, Duke University Medical Center, Durham, North Carolina, USA

## Abstract

Lipoteichoic acid (LTA) is a cell surface polymer of Gram-positive bacteria. LTA participates in host-microbe interactions including modulation of host immune reactions. It was previously reported that the major human pathogen *Streptococcus pneumoniae* and the closely related oral commensals *S. mitis* and *S. oralis* produce Type IV LTAs. Herein, using liquid chromatography/mass spectrometry (LC/MS)-based lipidomic analysis, we found that in addition to Type IV LTA biosynthetic precursors, *S. mitis*, *S. oralis*, and *S. pneumoniae* also produce glycerophosphate (Gro-P)-linked dihexosyl-diacylglycerol (DAG), which is a biosynthetic precursor of Type I LTA. Mutants in *cdsA* and *pgsA* produce dihexosyl-DAG but lack (Gro-P)-dihexosyl-DAG, indicating that the Gro-P moiety is derived from phosphatidylglycerol (PG), whose biosynthesis requires these genes. *S. mitis*, but neither *S. pneumoniae* nor *S. oralis*, encodes an ortholog of the PG-dependent Type I LTA synthase, *ltaS*. By heterologous expression analyses, we confirmed that *S. mitis ltaS* confers poly-(Gro-P) synthesis in both *Escherichia coli* and *Staphylococcus aureus*, and that *S. mitis ltaS* can rescue the severe growth defect of a *S. aureus ltaS* mutant. However, despite these observations, we do not detect a poly-(Gro-P) polymer in *S. mitis* using an anti-Type I LTA antibody. Moreover, (Gro-P)-linked dihexosyl-DAG is still synthesized by a *S. mitis ltaS* mutant, demonstrating that *S. mitis* LtaS does not catalyze the transfer of Gro-P from PG to dihexosyl-DAG. Finally, a *S. mitis ltaS* mutant has increased sensitivity to human serum, demonstrating that *ltaS* confers a beneficial but currently undefined function in *S. mitis*. Overall, our results demonstrate that *S. mitis*, *S. pneumoniae*, and *S. oralis* produce a (Gro-P)-linked glycolipid via a PG-dependent, *ltaS*-independent mechanism.

**Importance:** LTA is an important cell wall component synthesized by Gram-positive bacteria. Disruption of LTA production can confer severe physiological defects and attenuation of virulence. We report here the detection of a biosynthetic precursor of Type I LTA, in addition to the previously characterized Type IV LTA, in the total lipid extracts of *S. pneumoniae*, *S. oralis*, and *S. mitis*. Our results indicate that a novel mechanism is responsible for producing the Type I LTA intermediate. Our results are significant because they identify a novel feature of *S. pneumoniae*, *S. oralis*, and *S. mitis* glycolipid biology.

## Introduction

The Gram-positive bacteria *Streptococcus mitis* and *S. oralis*, members of the mitis group streptococci, are among the major oral colonizers that protect against human gingivitis via production of hydrogen peroxide, neutralization of acids, and secretion of antimicrobial compounds (1–5). They are also opportunistic pathogens that are among the leading causes of community-acquired bacteremia and infective endocarditis (IE) (6–8). Our understanding of how these organisms colonize, survive, and interact with the human host in these different niches is incomplete and requires further mechanistic study.

*S. pneumoniae* also belongs to the mitis group streptococci and shares > 99% identity in 16S rRNA sequence with both *S. mitis* and *S. oralis* (9, 10). *S. pneumoniae* mainly colonizes the mucosal surfaces of the human upper respiratory tract and is a well-known human pathogen causing pneumonia, meningitis, and otitis media, among other infections, and is a significant cause of morbidity and mortality worldwide (11, 12). Though *S. mitis*, *S. oralis*, and *S. pneumoniae* differ in their colonization abilities and pathogenic potential, multiple studies have shown that they share some common mechanisms of host-microbe interactions. For instance, *S. mitis* and *S. oralis* may serve as reservoirs of pneumococcal virulence-associated and antibiotic resistance genes (13–15); and immunity against *S. mitis* provides protection against *S. pneumoniae* colonization (16). We recently reported that *S. mitis*, *S. oralis*, and *S. pneumoniae* scavenge intermediates of human phospholipid metabolism and utilize them to synthesize the zwitterionic phospholipid phosphatidylcholine (PC), a pathway that potentially modulates human host immune responses (17, 18).

In addition to membrane phospholipids, another Gram-positive cell wall component that plays critical roles in host-microbe interactions is the lipoteichoic acid (LTA). LTA is a membrane lipid-anchored polymer typically consisting of either glycerophosphate (Gro-P) or ribitol-phosphate (Rbo-P) repeating units (19). LTAs with different chemical structures can trigger different immune responses from the host (20–22). According to their structural differences, LTAs have been grouped into five different types, among which the LTAs produced by *Staphylococcus aureus* (Type I) and *S. pneumoniae* (Type IV) have been extensively studied (23). Pneumococcal LTA was originally identified in 1943, and was named as F-antigen at that time due to its ability to cross-react with the Forssman antigen series (24). Its repeating unit consists of residues of 2-acetamido-4-amino-2,4,6-trideoxy-D-galactose (AATGal), D-glucose, Rbo-P, N-acetyl-D-galactosamine (GalNAc), and phosphocholine (25). Genes involved in the production of Type IV LTA were summarized by Denapaite *et al*. based on genomic predictions and previous experimental studies (26). Orthologs of these genes are also present in *S. oralis* and *S. mitis* genomes, except that for most *S. mitis* and *S. oralis* strains, the glucose glycosyltransferase is substituted with a galactose glycosyltransferase (26, 27). Structural analysis of the Type IV LTA produced by *S. oralis* strain Uo5 has confirmed the replacement of glucose residues by galactose, as well as revealed other differences relative to pneumococcal LTA in the repeating unit and branching structures (28).

*S. mitis* is the primary focus of the work presented here. The chemical structures of *S. mitis* LTAs vary among different strains, and conflicting data on *S. mitis* LTAs have been reported. Bergström *et al.* found that 39 of 77 *S. mitis* strains produce polysaccharide polymers detectable by monoclonal antibodies that separately target the pneumococcal Type IV LTA polymer backbone and phosphocholine residues (29). Among the remaining strains, some of them lack phosphocholine, such as *S. mitis* SK598, which produces a pneumococcal LTA-like polymer with the choline residues being replaced by ethanolamine (29, 30), while some might produce LTA of a different type. For example, another teichoic acid-like polymer consisting of repeating units of heptasaccharide phosphate was identified in cell lysates of *S. mitis* SK137 (29); however, whether this polymer is anchored to the membrane or the peptidoglycan is unknown. A few studies have reported detection of Type I-like LTA, a Gro-P polymer, from *S. mitis* clinical isolates using anti-Type I LTA antibodies (31–33). However, since these reports, species definitions among mitis group streptococci have been refined. A more recent reanalysis using the same detection technique did not detect Type I LTA in four *S. mitis* strains, including the type strain *S. mitis* ATCC 49456 (34). However, genomic analysis supports the possibility of Type I LTA synthesis in *S. mitis*, as *S. mitis* encodes an ortholog of the *S. aureus* type I LTA synthase gene, *ltaS* (26, 35). LtaS catalyzes the transfer of Gro-P from the membrane phospholipid phosphatidylglycerol (PG) and polymerizes the Gro-P units on a glycolipid anchor, forming Type I LTA (36, 37).

The goal of our study was to determine whether *S. mitis* produces multiple types of LTAs, and whether *S. mitis ltaS* mediates production of Type I LTA, using the type strain ATCC 49456 as a model. We used normal-phase liquid chromatography (NPLC)-electrospray ionization/mass spectrometry (ESI/MS) to analyze membrane lipids in the mitis group streptococci. This technique allows for analysis of LTA anchors and other LTA biosynthetic intermediates. We identified intermediates of Type IV LTA synthesis in *S. mitis*, *S. oralis*, and *S. pneumoniae*. To our surprise, a Type I-like LTA intermediate was observed not only in *S. mitis*, which encodes *ltaS*, but also in *S. oralis* and *S. pneumoniae*, which lack *ltaS* orthologs. Moreover, while *S. mitis* ATCC 49456 *ltaS* confers poly-(Gro-P) synthesis when heterologously expressed in *Escherichia coli* and a *S. aureus ltaS*-deficient mutant, we confirm that *S. mitis* ATCC 49456 does not produce a polymer detectable by a Type I LTA antibody. Importantly, *ltaS* contributes to *S. mitis* ATCC 49456 fitness, because deletion of *ltaS* impacted growth in human serum-supplemented medium. Overall, our results demonstrate that *S. mitis*, *S. oralis*, and *S. pneumoniae* synthesize intermediates of two structurally distinct lipid-anchored polymers, one Type IV LTA, and one a (Gro-P)-containing polymer whose full structure remains to be determined.

## Results

### Mitis group streptococci produce glycolipid intermediates of two structurally distinct LTAs

LTA is usually anchored to the membrane by a saccharide-linked diacylglycerol (DAG) glycolipid (23). Structure of the glycolipid anchor varies among different LTA types, bacterial species, and even culture conditions (38). In *S. pneumoniae*, the pseudopentasaccharide repeating units of Type IV LTA are proposed to be assembled on an undecaprenyl pyrophosphate (C_55_-PP) anchor, and then transferred to a glucosyl-DAG (Glc-DAG) anchor (Fig 1A) (25). In *S. aureus*, Type I LTA is typically assembled on a diglucosyl-DAG (Glc_2_-DAG) anchor (Fig 1A) (39).

**Fig. 1:**
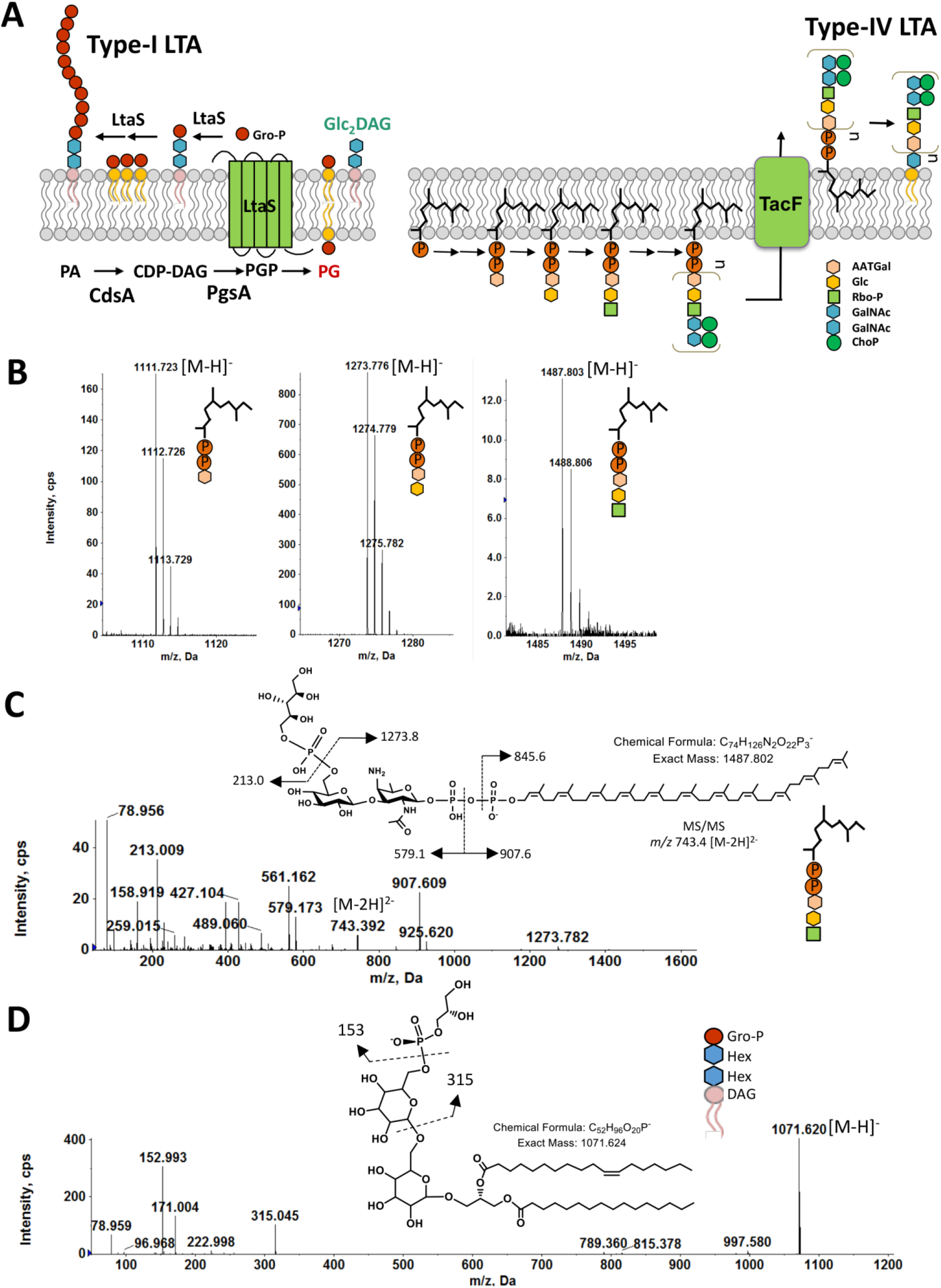
Detection of Type-IV LTA biosynthetic precursors and (Gro-P)-dihexosyl-DAG from the lipid extracts of *S. mitis* ATCC 49456 (SM61). Total lipids were extracted from *S. mitis* grown to mid-log phase in Todd Hewitt broth. A) Schematic of biosynthesis of *S. aureus* Type I and *S. pneumoniae* Type IV LTAs. B) Negative ion ESI mass spectra showing the [M-H]^−^ ions of C_55_-PP-AATGal, C_55_-PP-AATGal-Gal, and C_55_-PP-AATGal-Gal-(Rbo-P). These C_55_-PP-linked saccharides are intermediates involved in assembling the pseudopentasaccharide repeating units of Type IV LTA. C) MS/MS product ion mass spectrum of the *m/z* 743.4 [M-2H]^2−^ ion of C_55_-PP-AATGal-Gal-(Rbo-P) and the MS/MS fragmentation scheme. D) MS/MS of the *m/z* 1071.6 [M-H]^−^ ion of (Gro-P)-dihexosyl-DAG and the proposed fragmentation scheme. Abbreviations: PA, phosphatidic acid; CDP, cytidine diphosphate; PG, phosphatidylglycerol; PGP, PG-3-phosphate; Glc, glucose; C_55_-PP, undecaprenyl pyrophosphate; DAG, diacylglycerol; Gal, galacosyl; Gro-P, glycerophosphate; Rbo-P, ribitol-phosphate; AATGal, 2-acetamido-4-amino-2,4,6-trideoxy-D-galactose; GalNAc, N-acetyl-D-galactosamine; ChoP, phosphocholine; Hex, hexose.

*Listeria monocytogenes* also produces Type I LTA, which is linked to a galactosyl-glucosyl-DAG (Gal-Glc-DAG) anchor (40). Thus, lipid profiling has the potential to identify LTA intermediates, thereby revealing possible types of LTAs produced by a bacterium. To perform lipidomic analysis of mitis group streptococci, total lipids were extracted from bacterial cultures with a modified acidic Bligh-Dyer method and analyzed with NPLC-ESI/MS (41). We analyzed the type strain of *S. mitis* (ATCC 49456, referred to as SM61 hereafter), *S. oralis* (ATCC 35037), two clinically isolated *S. pneumoniae* strains (D39 and TIGR4), and *Streptococcus* sp. 1643 (referred to as SM43 hereafter), a human endocarditis isolate that was clinically identified as *S. mitis* but shares higher genomic identity with *S. oralis* (Table 1) (18, 42).

**Table 1:**
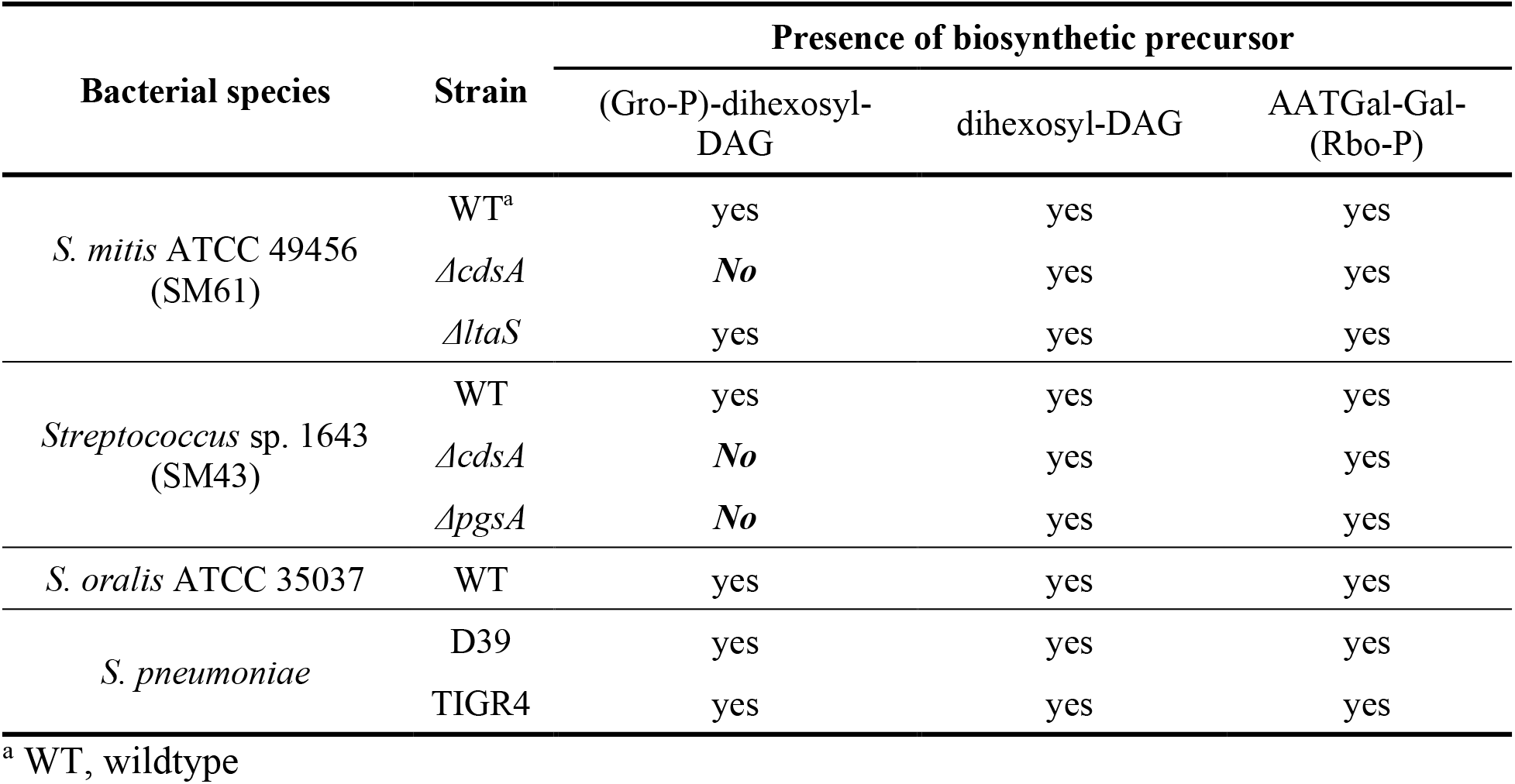
Detection of lipoteichoic acid intermediates from selected strains of mitis group streptococci

| Bacterial species | Strain | Presence of biosynthetic precursor |  |  |
| --- | --- | --- | --- | --- |
|  |  | (Gro-P)-dihexosyl-DAG | dihexosyl-DAG | AATGal-Gal-(Rbo-P) |
| *S. mitis* ATCC 49456 (SM61) | WT<sup>a</sup> | yes | yes | yes |
|  | *ΔcdsA* | **No** | yes | yes |
|  | *ΔltaS* | yes | yes | yes |
| *Streptococcus* sp. 1643 (SM43) | WT | yes | yes | yes |
|  | *ΔcdsA* | **No** | yes | yes |
|  | *ΔpgsA* | **No** | yes | yes |
| *S. oralis* ATCC 35037 | WT | yes | yes | yes |
| *S. pneumoniae* | D39 | yes | yes | yes |
|  | TIGR4 | yes | yes | yes |

<sup>a</sup> WT, wildtype

Three C_55_-PP-linked intermediates of Type IV LTA biosynthesis were detected in all strains analyzed. Specifically, these intermediates are C_55_-PP-linked AATGal ([M-H]^−^ at *m/z* 1111.7 of Fig. 1B, left), C_55_-PP-AATGal-Gal ([M-H]^−^ at *m/z* 1273.7 of Fig. 1B, middle), and C_55_-PP-AATGal-Gal-(Rbo-P) ([M-H]^−^ at *m/z* 1487.7 of Fig. 1B, right). Identifications of these species are supported by the exact mass measurement and tandem mass spectrometry (MS/MS). For example, Fig. 1C shows MS/MS of the doubly deprotonated [M-2H]^2−^ ion at *m/z* 743.4 for C_55_-PP-AATGal-Gal-(Rbo-P) along with the fragmentation scheme. In addition, we also detected (Gro-P)-dihexosyl-DAG ([M-H]^−^ at *m/z* 1071.6 of Fig. 1D), an intermediate that would be expected for Type I LTA. The exact mass measurement (*m/z* 1071.620) is consistent with the calculated [M-H]^−^ ion mass (*m/z* 1071.624) of (Gro-P)-dihexosyl-DAG containing C16:0 and C18:1 acyl chains. Furthermore, MS/MS of [M-H]^−^ ion at *m/z* 1071.6 for (Gro-P)-dihexosyl-DAG (16:0/18:1) along with the fragmentation scheme are shown in Fig 1D. The stereochemistry of the two hexoses cannot be discerned by MS/MS.

To confirm the possible monosaccharide identity of the DAG-linked sugars, *in silico* analyses were performed to identify orthologs of known glycolipid biosynthetic genes in the genomes of the tested strains. *S. pneumoniae* produces the glycolipid Gal-Glc-DAG (43), for which the biosynthetic genes have been partially identified. These genes can be separated into two major groups corresponding to the biosynthetic steps they are responsible for: 1) production of nucleotide-activated sugars, and 2) transferring of the activated sugar moieties to DAG (38). As shown in Table 2, these genes include: confirmed uridine diphosphate glucose (UDP-Glc) production gene *pgm* (encoding α-phosphoglucomutase) and *galU* (encoding UTP:α-glucose-1-phosphate uridyltransferase) (44); Leloir pathway genes that are proposed to produce uridine diphosphate galactose (UDP-Gal), specifically *galK* (encoding galactokinase) and *galT2* (encoding galactose-1-phosphate uridylyltransferase 2) (45, 46); and glycosyltransferases encoded by genes Spr0982 and *cpoA* which sequentially transfer Glc and Gal residues to DAG, respectively (47, 48). *S. pneumoniae* R6 is an avirulent and unencapsulated derivative of *S. pneumoniae* D39 (49). These two strains share the same glycolipid biosynthetic genes. Using *S. pneumoniae* R6 as reference, orthologs of Gal-Glc-DAG biosynthetic genes with ≥ 87% amino acid identity were identified in the genomes of SM61, *S. oralis* ATCC 35037, SM43, and *S. pneumoniae* TIGR4 (Table 2). This analysis suggests that the dihexosyl-DAG detected in our experiments is likely to be Gal-Glc-DAG.

**Table 2:**
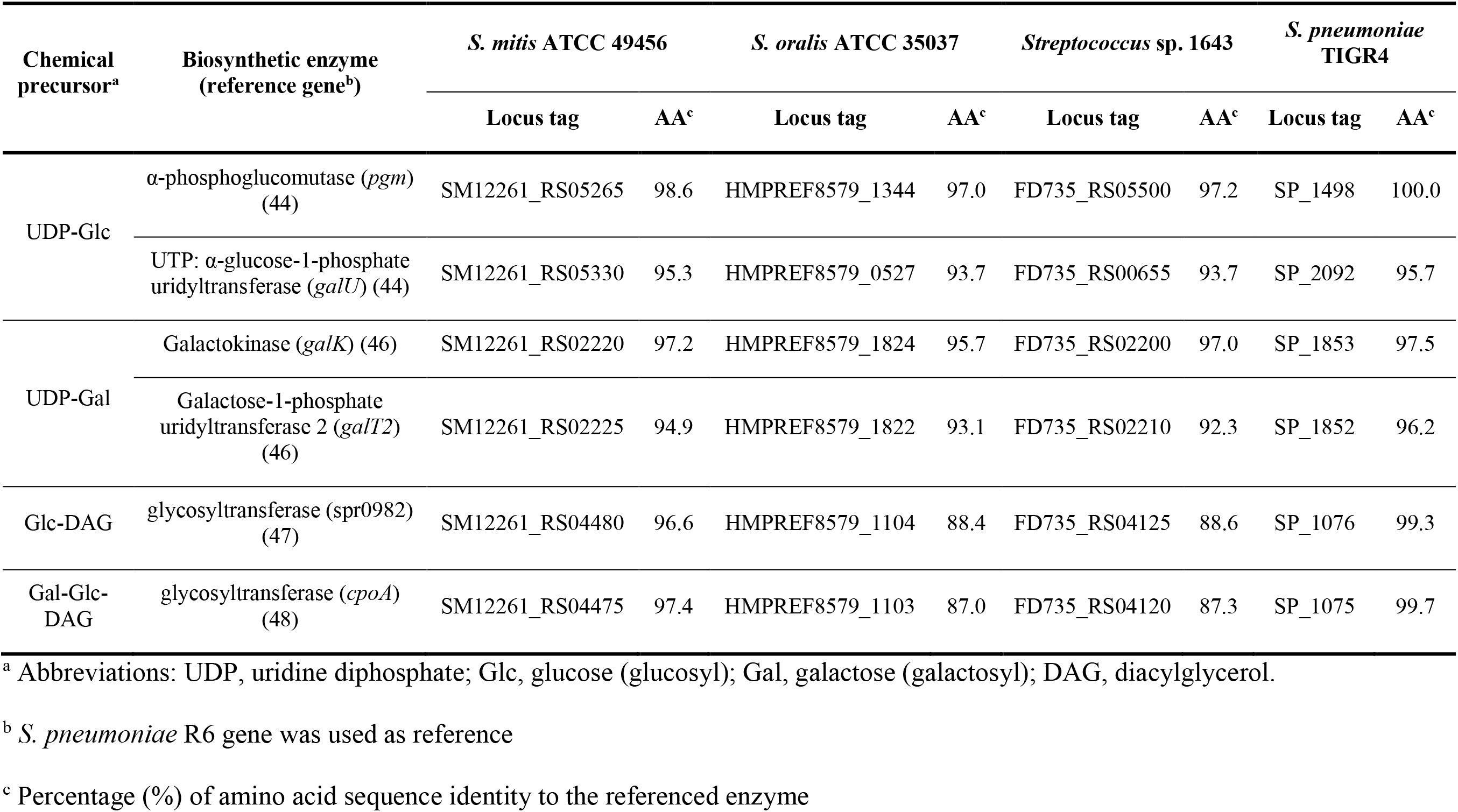
Orthologs of glycolipid biosynthetic genes

| Chemical precursor<sup>a</sup> | Biosynthetic enzyme (reference gene<sup>b</sup>) | *S. mitis* ATCC 49456 |  | *S. oralis* ATCC 35037 |  | *Streptococcus* sp. 1643 |  | *S. pneumoniae* TIGR4 |  |
| --- | --- | --- | --- | --- | --- | --- | --- | --- | --- |
|  |  | Locus tag | AA<sup>c</sup> | Locus tag | AA<sup>c</sup> | Locus tag | AA<sup>c</sup> | Locus tag | AA<sup>c</sup> |
| UDP-Glc | α-phosphoglucomutase (*pgm*) (44) | SM12261_RS05265 | 98.6 | HMPREF8579_1344 | 97.0 | FD735_RS05500 | 97.2 | SP_1498 | 100.0 |
| | UTP: α-glucose-1-phosphate uridylyltransferase (*galU*) (44) | SM12261_RS05330 | 95.3 | HMPREF8579_0527 | 93.7 | FD735_RS00655 | 93.7 | SP_2092 | 95.7 |
| UDP-Gal | Galactokinase (*galK*) (46) | SM12261_RS02220 | 97.2 | HMPREF8579_1824 | 95.7 | FD735_RS02200 | 97.0 | SP_1853 | 97.5 |
|  | Galactose-1-phosphate uridylyltransferase 2 (*galT2*) (46) | SM12261_RS02225 | 94.9 | HMPREF8579_1822 | 93.1 | FD735_RS02210 | 92.3 | SP_1852 | 96.2 |
| Glc-DAG | glycosyltransferase (spr0982) (47) | SM12261_RS04480 | 96.6 | HMPREF8579_1104 | 88.4 | FD735_RS04125 | 88.6 | SP_1076 | 99.3 |
| Gal-Glc-DAG | glycosyltransferase (*cpoA*) (48) | SM12261_RS04475 | 97.4 | HMPREF8579_1103 | 87.0 | FD735_RS04120 | 87.3 | SP_1075 | 99.7 |

<sup>a</sup> Abbreviations: UDP, uridine diphosphate; Glc, glucose (glucosyl); Gal, galactose (galactosyl); DAG, diacylglycerol.
<sup>b</sup> *S. pneumoniae* R6 gene was used as reference
<sup>c</sup> Percentage (%) of amino acid sequence identity to the referenced enzyme

### Biosynthesis of (Gro-P)-dihexosyl-DAG requires phosphatidylglycerol in mitis group streptococci

In *S. aureus*, the Gro-P of Type I LTA is produced from hydrolyzation of membrane PG (36), a process that is also required for Gro-P modification of streptococcal rhamnose-containing cell wall polysaccharides (50). To verify whether PG is the source of Gro-P for (Gro-P)-dihexosyl-DAG biosynthesis in mitis group streptococci, we analyzed the lipid profiles of *cdsA* and *pgsA* mutants. The gene *cdsA* is required for the synthesis of CDP-DAG, which is then converted by PgsA to produce phosphatidylglycerophosphate (PGP), the immediate precursor of PG (Fig. 1A) (18, 41). We previously reported that *cdsA* deletion mutants of *S. mitis* and *S. oralis* do not synthesize PG, nor does a *pgsA* deletion mutant of SM43 (18, 41) (Fig. 2). Thus, lipid anchor profiles of SM43 *cdsA* and *pgsA* deletion mutants were analyzed. While the dihexosyl-DAG glycolipid anchor (such as [M+Cl]^−^ at *m/z* 953.6 of Fig. 2) is observed in the wild type, *ΔcdsA*, and *ΔpgsA* strains, the (Gro-P)-linked dihexosyl-DAG (such as [M-H]^−^ at *m/z* 1071.6 of Fig. 2) is missing from the *ΔcdsA* and *ΔpgsA* strains. Identical anchor profiles were observed for the SM61 *cdsA* mutant (Table 1). These results demonstrate that *cdsA* and *pgsA*, or more specifically the ability to synthesize PG, are required for the biosynthesis of (Gro-P)-dihexosyl-DAG in SM61 and SM43.

**Fig. 2:**
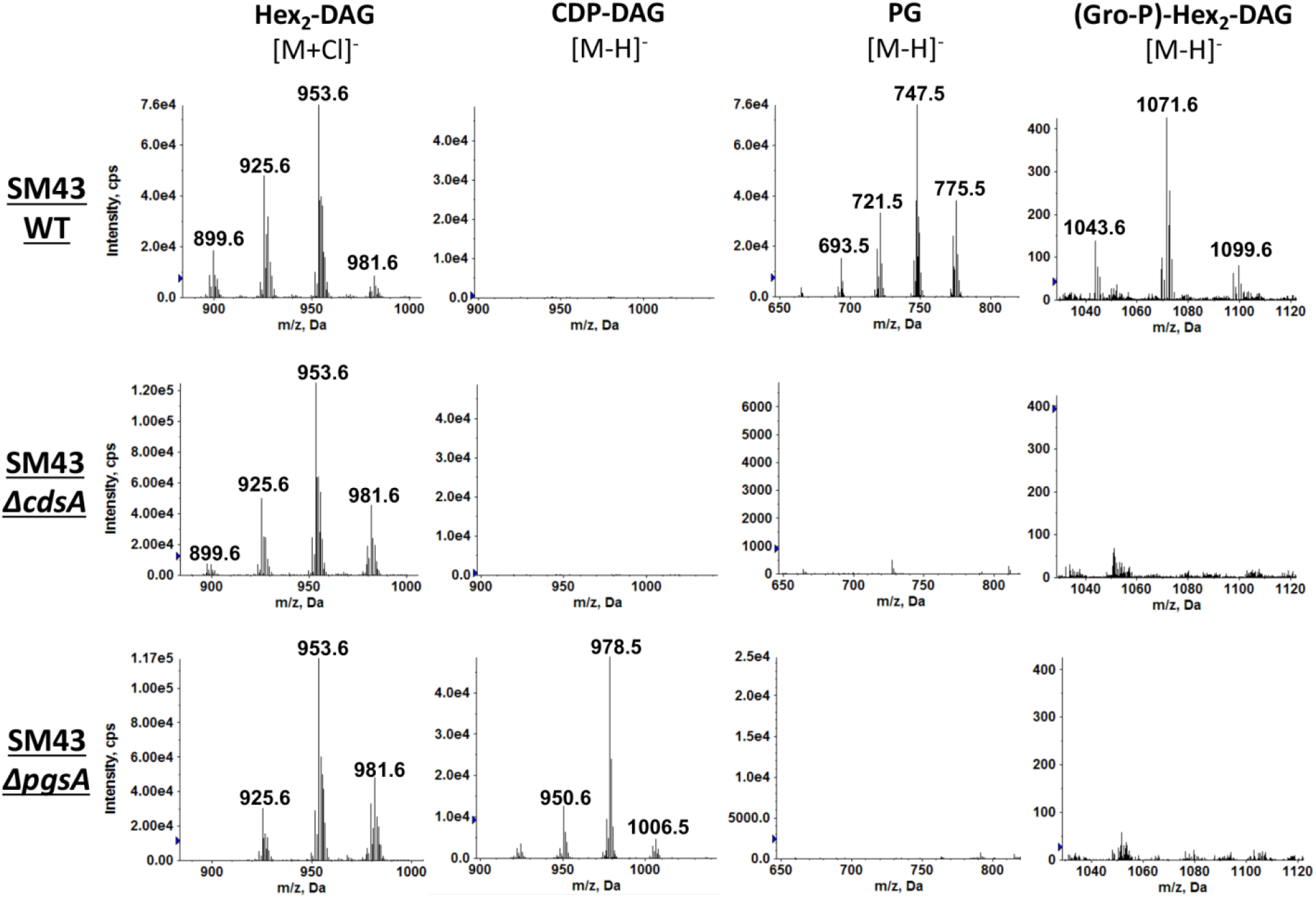
Negative ion ESI mass spectra showing the detection of phospholipids and anchor profiles from lipid extracts of *Streptococcus* sp. 1643 (SM43) wild type (WT), *ΔcdsA*, and *ΔpgsA* strains. Total lipids were extracted from SM43 cells grown to mid-log phase in Todd Hewitt medium. From left to right, each column correspondingly shows the mass spectra of the [M+Cl]^−^ ions of dihexosyl-diacylglycerol (Hex_2_-DAG) (retention time: ~8.0-10.0 min; most abundant *m/z* 953.6 for Hex_2_-DAG(16:0/18:1)), [M-H]^−^ ions of cytidine diphosphate-DAG (CDP-DAG) (retention time: ~21.5-22.5 min; most abundant *m/z* 978.5 for CDP-DAG(16:0/18:1)), phosphatidylglycerol (PG) (retention time: ~12.5-13.5 min; most abundant *m/z* 747.5 for PG (16:0/18:1)), and glycerophosphate (Gro-P) linked Hex_2_-DAG (retention time: ~20.0-20.5 min; most abundant *m/z* 1071.6 for (Gro-P)-Hex_2_-DAG(16:0/18:1)). The identification of these lipid species is supported by both exact mass measurement and MS/MS.

### S. mitis, S. oralis, and S. pneumoniae cell extracts do not react with a Type I LTA antibody

Currently, enzymes known to transfer Gro-P from PG for Gro-P polymer synthesis or Gro-P modification include 1) *S. aureus* LtaS, the Type I LTA synthase that produces poly-(Gro-P) (36), 2) *L. monocytogenes* LtaP, the Type I LTA primase that has a very similar overall structure and active site sequences with LtaS, except links only the first Gro-P unit to the glycolipid anchor (35, 40), and 3) the recently identified streptococcal Gro-P transferase GacH that links Gro-P to cell wall-attached glycopolymers (50). Bioinformatic analyses predict no orthologs of either *ltaP* or *gacH* in the genomes of the mitis group streptococci assessed here, yet an ortholog of *ltaS* is present in *S. mitis*, as previously reported (35).

If *S. mitis ltaS* functions the same as its ortholog in Type I LTA-producing bacteria like *S. aureus*, polymers of Gro-P will be produced and may be detectable using an anti-Type I LTA antibody. Western blot analysis using a previously described anti-Type I LTA antibody was conducted for SM61, SM43, *S. oralis* ATCC 35037 and *S. pneumoniae* strains. No signal was detected from cell lysates of these strains (Fig. 3), nor from cell lysates of SM61 that over-expresses *ltaS in trans* from an anhydrotetracycline-inducible vector (Fig. S1). These results are in accordance with previous observations of no immunoluminescent detection of Gro-P polymers in SM61 (34). The validity of the antibody was confirmed by positive signals detected from cell lysates of *S. agalactiae*, *S. pyogenes*, and *S. aureus*, all three of which produce Type I LTA (Fig. 3) (36, 51, 52). Interestingly, no signal was detected from cell lysate of *Enterococcus faecalis* OG1RF (Fig. 3), another bacterium known to produce Type I LTA (53, 54), which, as reported previously, is poorly recognized by the anti-Type I LTA antibody (55).

**Fig. 3:**
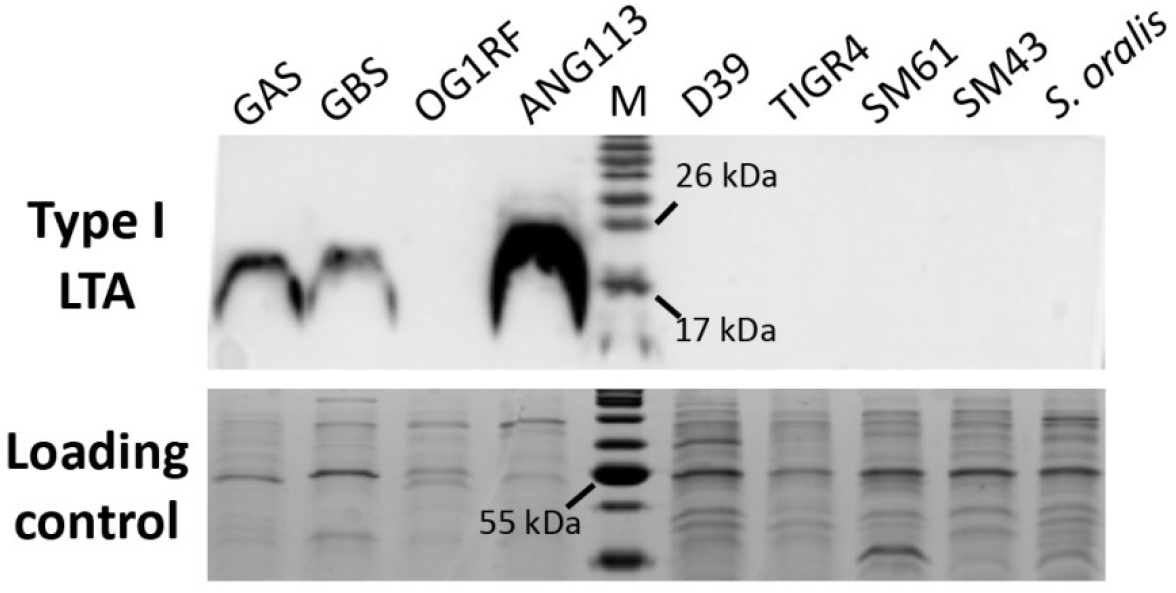
Detection of Type I LTA. Cell lysates from over-night cultures of *Streptococcus pyogenes* NZ131 (GAS), *S. agalactiae* A909 (GBS), *Enterococcus faecalis* OG1RF (OG1RF), *Staphylococcus aureus* (ANG113), *S. pneumoniae* D39 (D39), *S. pneumoniae* TIGR4 (TIGR4), *S. mitis* ATCC 49456 (SM61), *Streptococcus* sp. 1643 (SM43), and *S. oralis* ATCC 35037 (*S. oralis*) were analyzed. Anti-Type I LTA antibody was used to detect the production of Type I LTA. Loading control was stained with Commassie blue.

### S. mitis LtaS mediates production of poly-(Gro-P) in an E. coli heterologous host

For the following analyses, the *S. mitis* type strain ATCC 49456 (SM61) was used as a model, and its *ltaS* ortholog (SM12261_RS03435) was renamed *ltaS*. We heterologously expressed *S. mitis ltaS* in *E. coli* to verify the function of the gene. This approach was previously used in studies of *S. aureus ltaS* (36). Plasmid pET-ltaS (Table 3) was constructed so that the expression of *S. mitis ltaS* could be induced with IPTG in *E. coli*. As shown in Fig. 4A, with the addition of IPTG, detectable bands produced by anti-Type I LTA antibody targeting were observed for *E. coli* (pET-ltaS), demonstrating that *S. mitis ltaS* is sufficient to mediate the production of poly-(Gro-P).

**Table 3:**
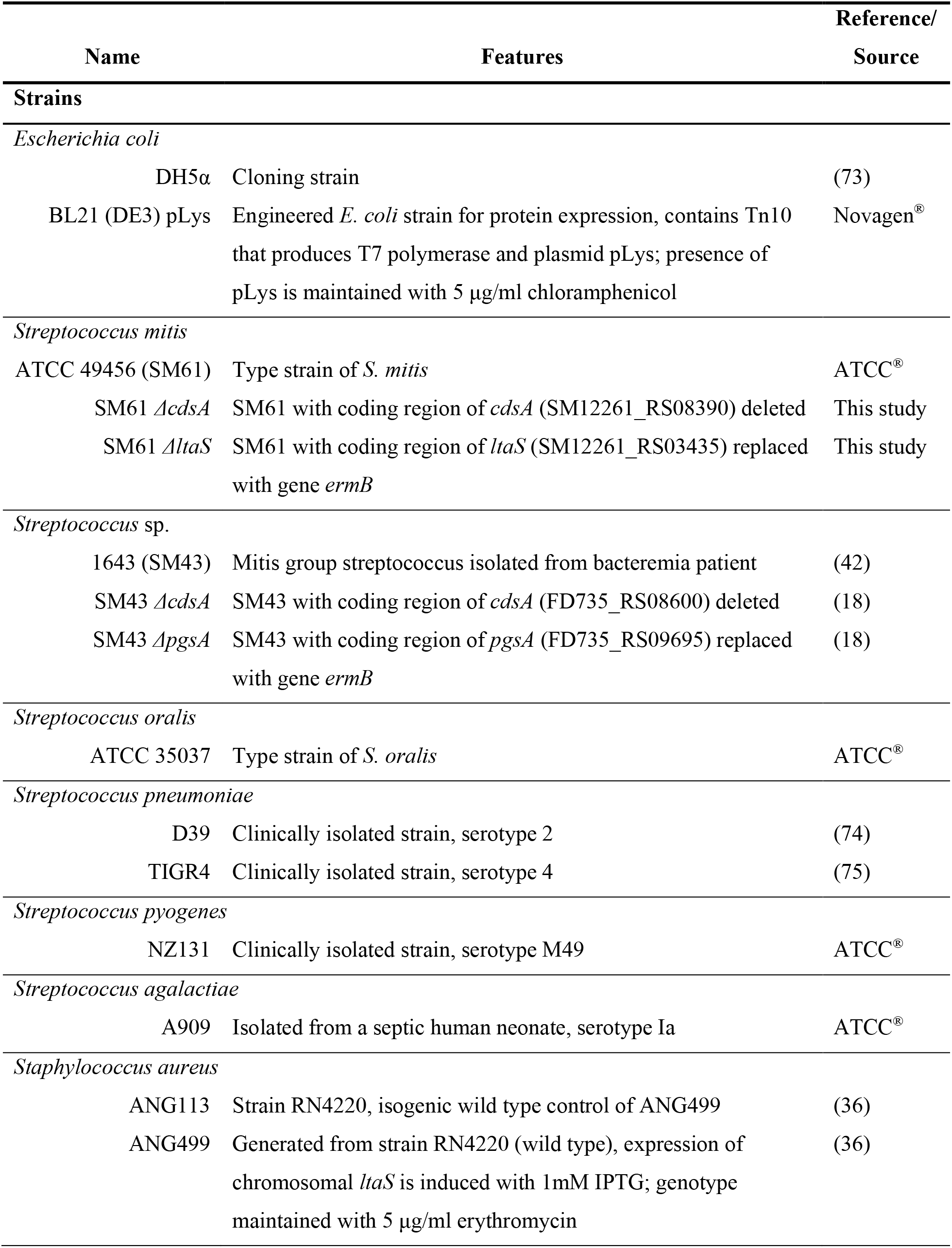
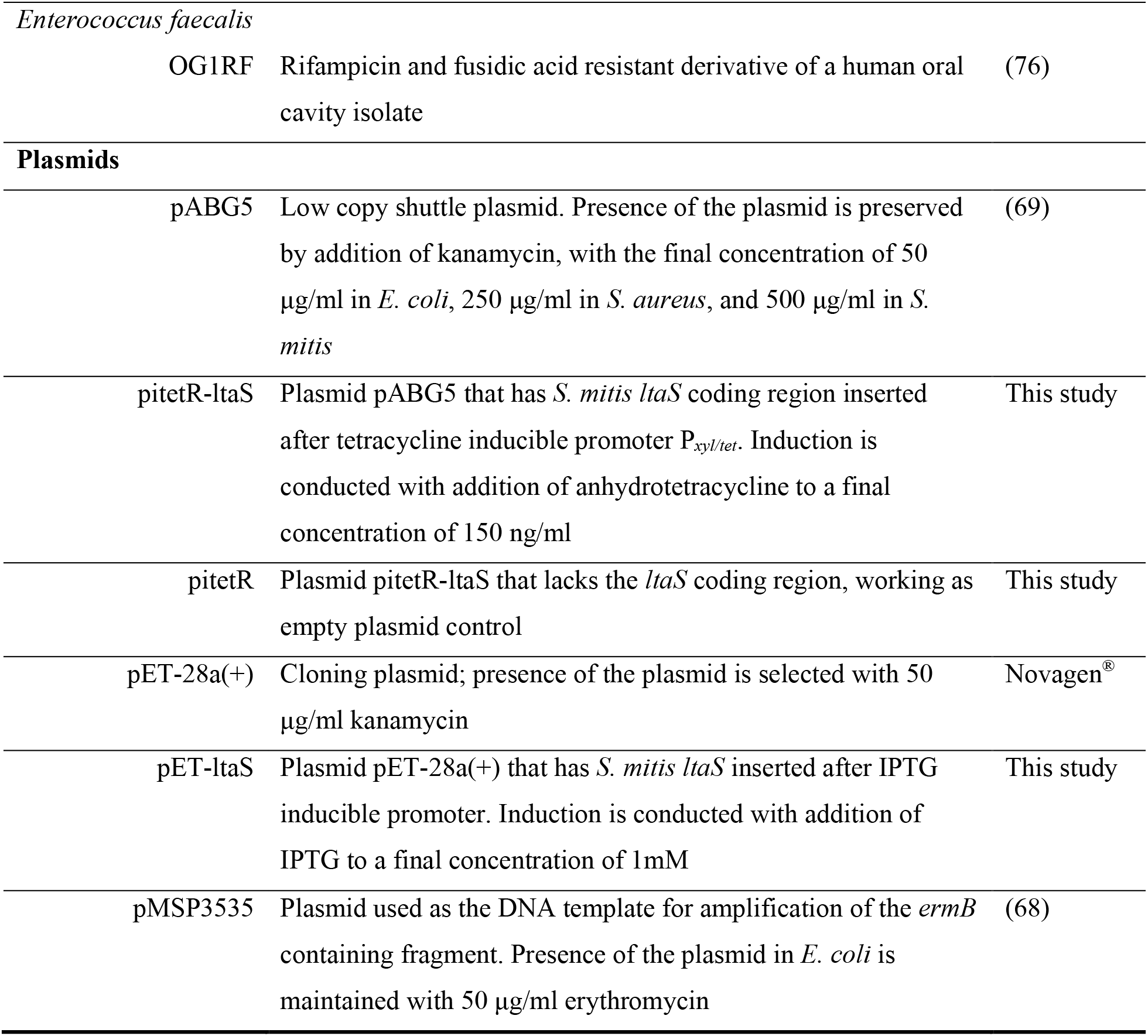
Bacterial strains and plasmids used in this research

**Fig. 4:**
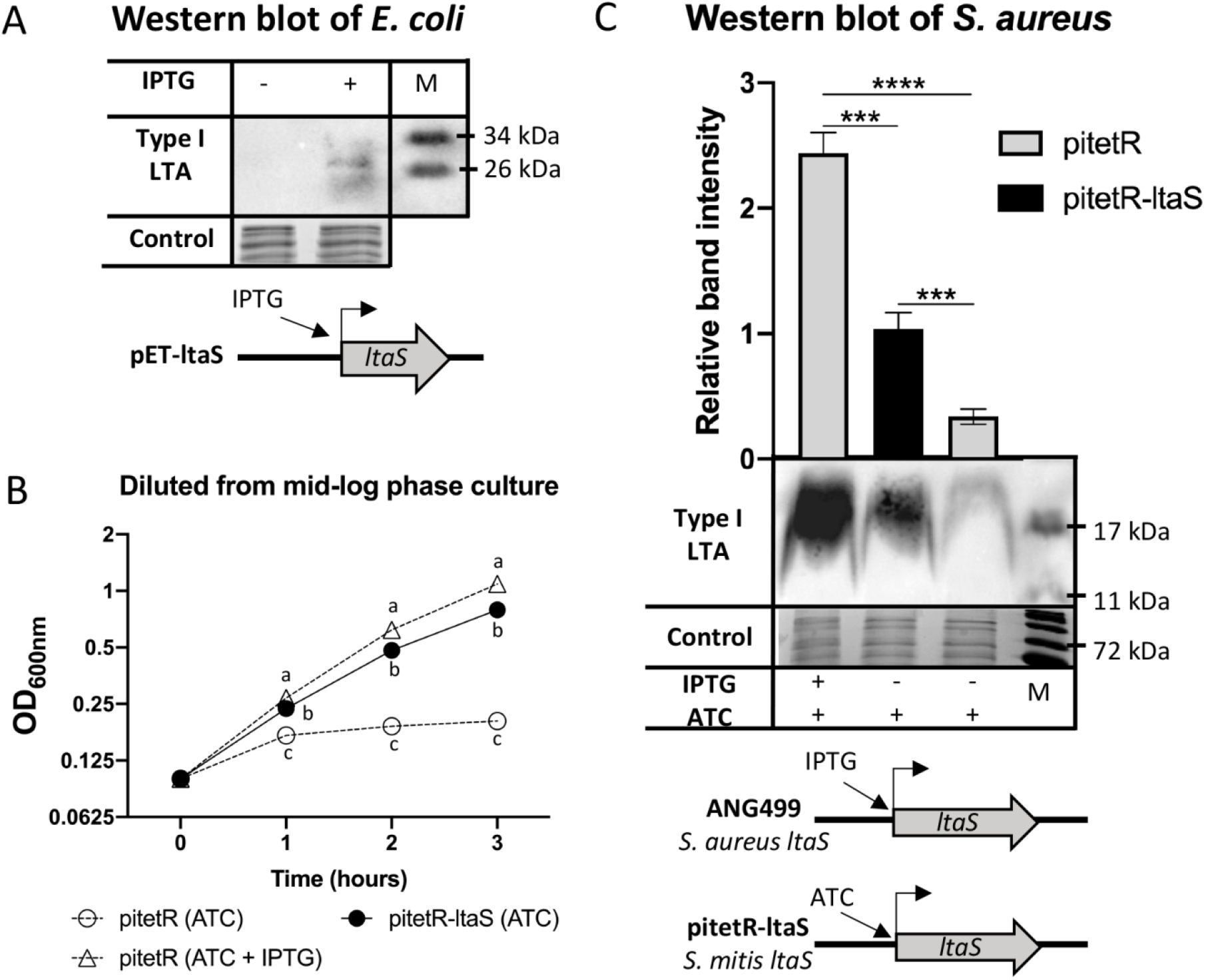
Heterologous expression of *S. mitis ltaS* in *E. coli* and *S. aureus*. A. Western blot detection of Gro-P polymers from *E. coli* containing plasmid pET-ltaS grown in liquid Luria-Bertani (LB) media with the addition of isopropyl β-D-1-thiogalactopyranoside (IPTG) and without IPTG. IPTG was added to mid-log phase bacterial cultures followed by another 30 minutes incubation at 37°C before cell pelleting. Three biological independent replicates were performed for each sample. B. Growth curves of *S. aureus* ANG499 containing either pitetR or pitetR-ltaS grown in Tryptic Soy Broth (TSB) with the addition of either 150 ng/ml anhydrotetracycline (ATC) only or 150 ng/ml ATC and 0.5 mM IPTG as indicated. Samples were grown in TSB with 0.5 mM IPTG over-night, followed by sub-culturing into fresh TSB with the indicated addition of induction reagents and incubated for 3 hours. Then, another sub-culturing to an OD_600nm_ of 0.1 with fresh media same as previous incubation was performed. After the second sub-culture, OD_600nm_ values were measured every hour and plotted. C. Western blot detection of Type I LTA from *S. aureus* ANG499 containing either pitetR or pitetR-ltaS. Samples were grown in the same way as described in B, after the first sub-culturing and incubation, cells equal to 1ml of OD_600nm_ at 1.2 were harvested followed by lysate preparation and immunodetection. Schematics of induction expression of chromosomal or plasmid carried *ltaS* were shown in both A & C. Loading controls of both A & C were stained with Commassie blue. Western blot band intensity in C was normalized to the loading control and the pitetR-ltaS sample. For B and C, 4 biological replicates were performed; averages of the sample values were plotted with the error bar stands for standard deviation. Statistical analyses were performed with one-way ANOVA; significant difference was determined by *P*-value < 0.05. For B, at a given time point, letter “a”, “b”, and “c” each represents a statistical group that is significantly different from other groups; *P*-values of all group comparisons are < 10^−6^. For C, “***” indicates 10^−5^ < *P*-value < 10^−6^; “****” indicates *P*-value < 10^−6^.

### S. mitis ltaS complements a S. aureus ltaS mutant for Type I LTA production

In *S. aureus*, LtaS is required for proper cell division and efficient cell growth at 37°C (36, 56). To further confirm the physiological function of *S. mitis ltaS* in Gram-positive cells, we expressed it in a previously reported *S. aureus* strain that has its native *ltaS* gene under the control of an IPTG-inducible promoter (strain ANG499). Without IPTG, ANG499 is deficient for Type I LTA production and has a growth defect when cultured at 37°C (36, 56). *S. mitis ltaS* was introduced into ANG499 by the plasmid pitetR-ltaS (Table 3), which has the *S. mitis ltaS* coding region under the control of the tetracycline-inducible promoter P_*xyl/tet*_. Addition of anhydrotetracycline (ATC) induces expression of *S. mitis ltaS*. Note that we included ATC in all experimental cultures described below, because we observed an ATC-dependent growth defect that confounded direct comparison of ATC+/ATC-cultures (Fig. S2).

As expected, ANG499 with the empty plasmid vector pitetR grew more slowly and reached a lower final OD_600nm_ value when cultured without IPTG as compared to with IPTG (Fig. 4B). As expected, Type I LTA production by *S. aureus* LtaS was induced by IPTG, confirmed by Western blot analysis (Fig. 4C) and detection of Type I LTA intermediates (Gro-P)2-Glc_2_-DAG ([M-H]^−^ ion at *m/z* 1214.6 of Fig. 5A) and alanine-linked (Gro-P)2-Glc_2_-DAG ([M-H]^−^ ion at *m/z* 1285.7 of Fig. S3). Strikingly, the growth of ANG499 was also rescued by the expression of *S. mitis ltaS* from pitetR-ltaS (Fig. 4B), and Type I LTA production was observed, as shown in Western blot (Fig. 4C) and lipidomic analysis (Fig. 5A & Fig. S3). These data demonstrate that *S. mitis ltaS* can complement the function of *S. aureus ltaS* and promote production of Type I LTA in *S. aureus*. Surprisingly, (Gro-P)-Glc_2_-DAG ([M-H]^−^ ion at *m/z* 1059.6 of Fig. 5B) was detected at comparable levels from all *S. aureus* cultures, including the natively *ltaS*-deficient strain in the absence of IPTG induction.

**Fig. 5:**
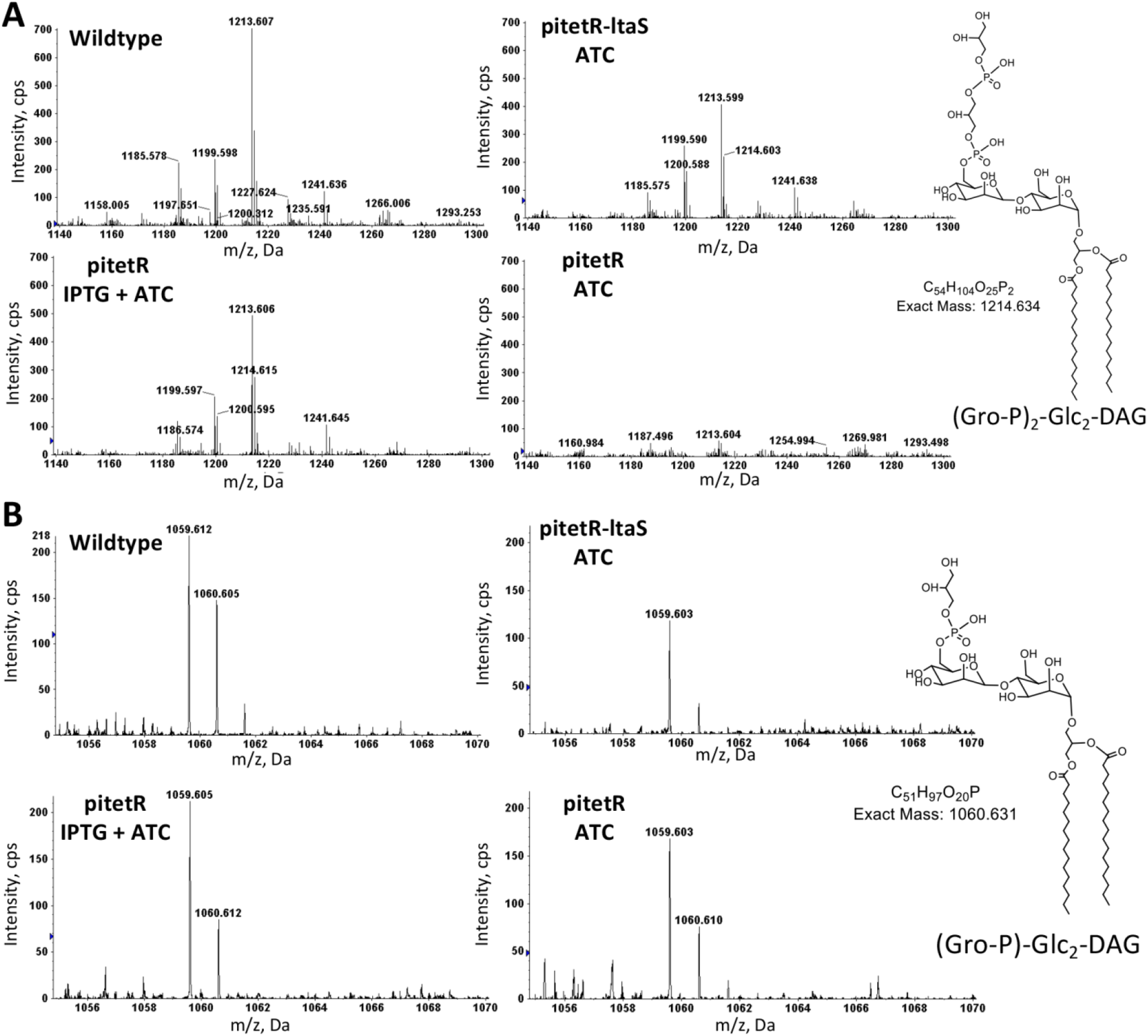
MS detection of Type I LTA biosynthetic precursors that contain one or two Gro-P units in the lipid extracts of *S. aureus*. *S. aureus* strain ANG113 (wildtype), ANG499 containing plasmid pitetR-ltaS (pitetR-ltaS), and ANG499 containing the vector control pitetR (pitetR) were grown in liquid Tryptic Soy medium to late exponential phase with the addition of ATC and IPTG as indicated. Total lipids were extracted and analyzed with NPLC-ESI/MS in the negative ion mode. Shown are the mass spectra of the deprotonated [M-H]^−^ ions for (Gro-P)-Glc_2_-DAG (retention time: ~20.0-20.5 min; most abundant *m/z* 1059.6) (A) and (Gro-P)_2_-Glc_2_-DAG (retention time: ~22.5-23.0 min; most abundant *m/z* 1213.6) (B). Abbreviations: Gro-P, glycerophosphate; Glc, glucosyl; DAG, diacylglycerol. Three biologically independent replicates were performed for each strain under each indicated culture condition.

### S. mitis lacking ltaS has increased serum susceptibility

To investigate functions of *ltaS* in *S. mitis*, *ltaS* was deleted and exchanged for the erythromycin resistance marker *ermB*, generating *S. mitis ΔltaS*. Of note, (Gro-P)-dihexosyl-DAG was still detected in the *S. mitis ΔltaS* strain, demonstrating that LtaS is not required for the addition of the Gro-P unit to the dihexosyl-DAG (Table 1).

Unlike *S. aureus*, which requires *ltaS* for efficient growth, deletion of *ltaS* in *S. mitis* does not confer a growth defect under laboratory culturing conditions. Specifically, when growing in Todd Hewitt Broth at 37°C, the doubling time of *ΔltaS* is 39.8 (± 3.7) minutes, which is not significantly different from the 40.2 (± 3.5) minute doubling time of wild type *S. mitis* (Fig. S4). Considering that the growth deficiency of *S. aureus* lacking *ltaS* could be mitigated by culturing at a lower temperature (56), the growth of *S. mitis* wild type and *ΔltaS* strains cultured at a higher temperature was measured, to determine whether the *ltaS* mutant was compromised for temperature-related stresses. The temperature 42°C was chosen as a representative of fever. Both wild type and *ΔltaS* strains exhibited slower growth at 42°C compared to 37°C; however, no significant difference in growth rate was observed between the strains (46.2 (± 3.0) and 47.4 (±3.8) minute doubling times for the wild type and *ΔltaS* strains, respectively). Moreover, no difference in susceptibilities to antibiotics targeting peptidoglycan biosynthesis, membrane integrity, and protein synthesis were observed (Table S1). Thus, under these laboratory culture conditions, *ltaS* is not essential for the growth of *S. mitis*.

In addition, a potential role for *ltaS* in host-microbe interactions was investigated. As an oral commensal, the environment *S. mitis* colonizes is exposed to human gingival crevicular fluid, which is an extrudant of serum with lower concentrations of complement (57). Moreover, when invading the bloodstream and causing bacteremia and infectious endocarditis, *S. mitis* is constantly exposed to blood. Thus, human serum is a useful medium component for laboratory reconstruction of the host growth conditions. Supplementation of human serum into chemically defined medium (CDM) promotes the growth of *S. mitis* compared to non-supplemented CDM (Fig. 6). Deletion of *ltaS* does not confer a significant difference in growth in Todd Hewitt broth or un-supplemented CDM; but does result in a significant growth deficiency in human serum-supplemented CDM, and makes *S. mitis* more sensitive to the killing effect of complete serum (Fig. 6). These results suggest that although *ltaS* is not required for growth of *S. mitis* under laboratory conditions, it is involved in interactions with human serum factors. Further investigation is needed to elucidate such interactions.

**Fig. 6:**
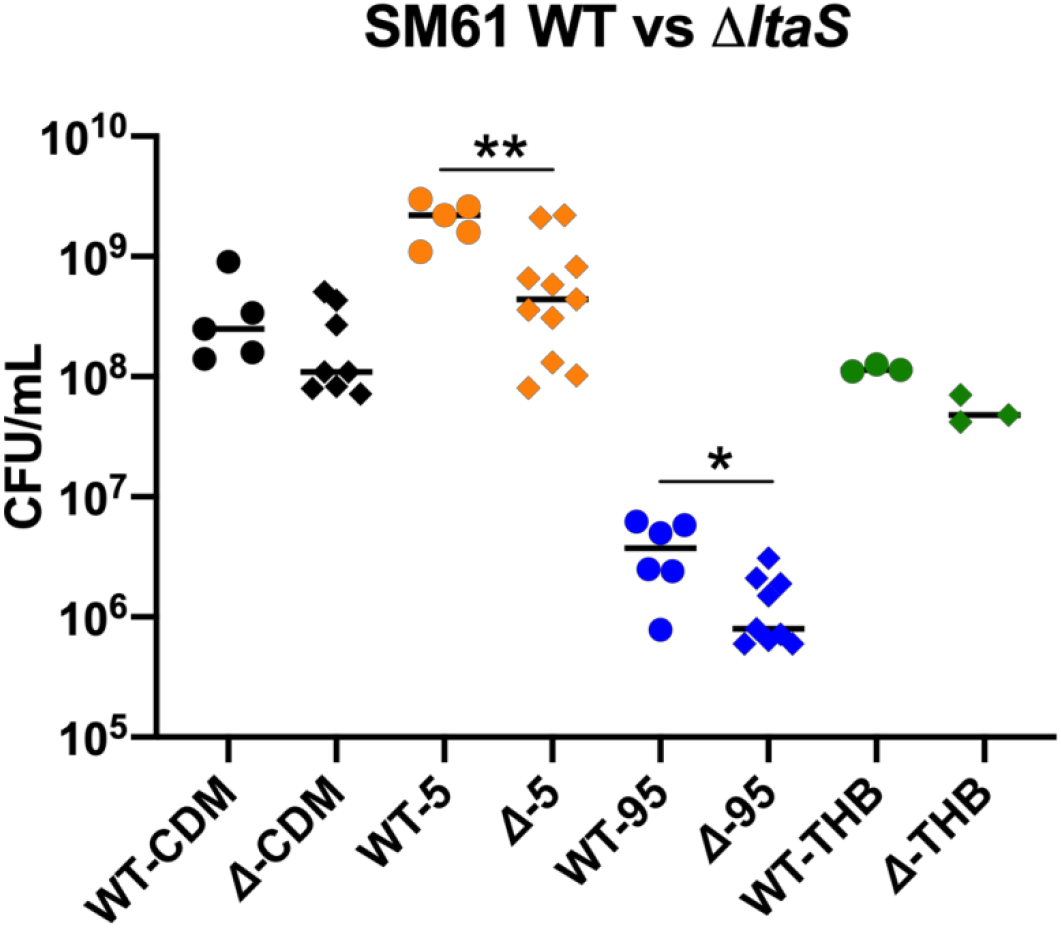
Deletion of *ltaS* alters the responses of *S. mitis* ATCC 49456 (SM61) to human serum. Wild type (WT) or *ΔltaS* SM61 (*Δ*) strains were cultured in chemically defined medium (CDM), CDM with 5% human serum (5), 95% human serum (95) with 5% phosphate buffered saline, and Todd Hewitt broth (THB). The CFU/mL of cultures after 8 hours incubation are shown. Each dot represents a biological independent repeat. Statistical analysis was performed with the Mann-Whitney method. Statistical significant is defined by *P*-value < 0.05, and indicated in the plot as “*” for 0.01 < *P*-value < 0.05, “**” for 0.001 < *P*-value < 0.01.

## Discussion

In this work, we used NPLC-ESI/MS to analyze the glycolipid profiles of *S. mitis*, *S. oralis*, and *S. pneumoniae* strains. For all of the tested strains, biosynthetic intermediates of two structurally different LTAs were detected (Fig. 1 and Table 1). Firstly, consistent with literature, the biosynthetic intermediates of the Type IV LTA were detected, which is in agreement with genomic analysis of the biosynthetic genes (26). The second distinct LTA is indicated by the detection of (Gro-P)-dihexosyl-DAG, which is similar to Type I LTA polymers and unexpected based on previous reports, and thus has been the focus of this study.

Based on genomic analysis, we proposed that the newly identified (Gro-P)-dihexosyl-DAG is structured as (Gro-P)-Gal-Glc-DAG. The glycolipid Gal-Glc-DAG has been reported as the dominant glycolipid species in *S. pneumoniae*, and our prediction is in accordance with this previous report (43). However, the full pathway for Gal-Glc-DAG synthesis has not been fully experimentally verified in the mitis group streptococci; the stereochemistry of the hexoses requires further confirmation with structural analysis, such as with NMR.

The PG-dependent (Gro-P)-dihexosyl-DAG biosynthetic process in *S. mitis* was then investigated, which led to the main focus of this study, functional verification of *S. mitis ltaS*. Through heterologous expression, we confirmed that *S. mitis ltaS* could directly synthesize Gro-P polymers in both *E. coli* and *S. aureus*. However, it appeared that *S. mitis* LtaS functions somewhat differently from *S. aureus* LtaS, as the expression of *S. mitis ltaS* does not fully complement the growth deficiency and the amount of Type I LTA produced (Fig. 4 B & C), which is not unexpected considering that *S. mitis* and *S. aureus* LtaS share only 38% sequence identity (26).

We did not detect a Gro-P polymer in wild type *S. mitis* using Western blot analysis. Explanations as to why we could not detect the polymer include: 1) *S. mitis* does not produce the Gro-P polymer; instead, (Gro-P)-dihexosyl-DAG is the complete and final product; 2) a very low amount of the Gro-P polymer is produced under the culture conditions investigated here; or 3) unique structural modifications on the Gro-P polymer hinder antibody recognition. Further large-scale purification and structural analysis of the (Gro-P)-dihexosyl-DAG-containing polymer produced by mitis group streptococci is required.

The findings that (Gro-P)-dihexosyl-DAG is still present in *S. mitis ΔltaS*, as well as in *S. oralis* and *S. pneumoniae*, which are species that encode no orthologs of *ltaS*, suggest the existence of an unknown PG-dependent Gro-P transferase in these species that is responsible for the synthesis of (Gro-P)-dihexosyl-DAG. Unexpectedly, (Gro-P)-Glc_2_-DAG is also seen in *S. aureus* deficient for *ltaS*, suggesting that an unidentified Gro-P biosynthetic enzyme(s) or biological process(es) may exist in *S. aureus* as well, but this is more speculative.

In other Gram-positive pathogens that synthesize Type I LTA, LtaS and its product, LTA, are essential for proper cell division (40, 56, 58, 59). Inhibiting the function of LtaS is effective in extending the survival of *S. aureus* infected mice (60) and sensitizing multi-drug resistant *E. faecium* to antibiotics (61). Though *S. mitis ltaS* is not essential for proper growth of the bacterium in normal laboratory media, nor for synthesizing (Gro-P)-dihexosyl-DAG, it does provide some advantage to *S. mitis* when human serum is present in the culture media.

In summary, we provide evidence that a Type I-like LTA might co-exist with Type IV LTA in *S. mitis*, *S. oralis*, and *S. pneumoniae*, and queried the role of *ltaS* in this process in a model *S. mitis* strain. To our knowledge, there is only one previous report which documents a bacterial species producing two structurally different LTAs, in *S. suis*, an invasive pathogen of pigs (62). Our lipidomic and genomic studies show that that we have an incomplete understanding of glycolipids and LTAs in mitis group streptococci, and their potential roles in host-microbe interactions.

## Materials and methods

### Bacterial strains and growth conditions

Unless indicated, *E. coli* were grown in Luria-Bertani (LB) medium, *Streptococcus* strains were grown in Todd Hewitt (TH) medium (BD Biosciences) with *S. pneumoniae* grown in TH medium supplemented with 0.5% yeast extract (BD Biosciences), and *E. faecalis* and *S. aureus* were grown in Tryptic Soy (TS) medium (BD Biosciences). All bacterial cultures were incubated at 37°C, unless otherwise noted. Streptococci were cultured with 5% CO_2_. Chemically defined medium (CDM) was made as previously described, with the addition of 0.5 mM choline (63). Human serum-supplemented medium was made through addition of complete human serum (Sigma Aldrich) into CDM to a final concentration of 5% (v/v). Bacterial strains and plasmids as well as the concentrations of antibiotics and expression-inducing reagents used in this research are listed in Table 3.

### Sequence analysis

Orthologs of glycolipid biosynthetic genes were identified through using the BLASTp function against the NCBI database (64). Specifically, genes of *S. pneumoniae* R6 (NC_003098.1) were used as reference. The encoded amino acid sequences were input into BLASTp to search against non-redundant protein database of *S. mitis* ATCC 49456 (taxid: 246201), *S. oralis* ATCC 35037 (taxid: 655813), *Strepcococcus* sp. 1643 (taxid: 2576376), and *S. pneumoniae* TIGR4 (taxid: 170187) individually. The *ltaS* (SM12261_RS03435) ortholog in *S. mitis* ATCC 49456 was identified similarly, with the amino acid sequence of *S. aureus* LtaS (SAV0719) (36) being the reference. Orthologs were determined by query coverage > 95% and E-value < 10^−120^.

### Mutant generation

Deletion of *cdsA* (SM12261_RS08390) in *S. mitis* ATCC 49456 was conducted as previously described (65–67). Briefly, approximately 2 kb flanking regions on either side of *cdsA* were amplified using Phusion polymerase (Thermo Fisher). PCR products were digested with restriction enzyme XmaI (New England Biolabs) and ligated with T4 DNA ligase (New England Biolabs). Ligated products were amplified using primers 61cdsA_Up_F and 61cdsA_Dwn_R (Table S2), followed by gel extraction with the QIAquick Gel Extraction Kit (Qiagen) per the manufacturer’s instruction. The linear construct was transformed into *S. mitis* via natural transformation as described previously (67). The *ΔcdsA* mutant was selected with 35 μg/ml daptomycin and confirmed with Sanger sequencing (Massachusetts General Hospital DNA Core) of the PCR product of the *cdsA* deletion region.

Deletion of *ltaS* in *S. mitis* ATCC 49456 was conducted similarly with some slight modifications. Specifically, a 1 kb DNA fragment containing *ermB* was generated through PCR amplification using plasmid pMSP3535 as the template (68). Then, splicing by overlap extension PCR was performed to produce a 5 kb amplicon that sequentially contained a 2 kb fragment upstream of *ltaS*, a 1 kb *ermB*-containing fragment in reverse orientation, and a 2 kb fragment downstream of *ltaS*. The PCR product was analyzed on a 0.8% agarose gel and extracted using the QIAquick Gel Extraction Kit (Qiagen) per the manufacturer’s instruction. Transformation of the 5 kb amplicon into *S. mitis* was performed as described previously (67). The *ΔltaS* mutant was selected with 20 μg/ml erythromycin and confirmed with Illumina genome sequencing (UTD Genome Core Facility).

### Plasmid construction

Plasmids used in this research are listed in Table 3 with description of their functions. All primers used in this research are listed in Table S2.

The shuttle plasmid pABG5 was used for heterologous gene expression in Gram-positive bacteria (69). Specifically, the DNA fragment containing the *S. mitis ltaS* coding region was amplified using primers LtaS_F and LtaS_R, and the pABG5 plasmid backbone was linearized through PCR using primers pABG5-5 and pABG5-3. Gibson assembly was conducted per the manufacturer’s instructions (NEBuilder HiFi DNA Assembly Master Mix, New England Biolabs), followed by transformation of the product into *E.coli* DH5α. The pABG5 with *ltaS* insert was further linearized with primers YW55 and YW56 and ligated with an 848 bp DNA fragment via Gibson assembly, producing the plasmid pitetR-ltaS. The 848 bp fragment contained a tetracycline-controlled promoter P_*xyl/tet*_ and the tetracycline repressor gene *tetR* in reverse orientation. Insertion of this 848 bp fragment immediately upstream of the *ltaS* coding region makes *ltaS* expression inducible by anhydrotetracycline (ATC) addition. Sequence of the 848 bp fragment was obtained from the Addgene sequence database (70), and the fragment was synthesized commercially (Integrated DNA Technologies). Induced production of the target gene *ltaS* was confirmed with Western blot. The empty vector control pitetR was constructed via linearization of pitetR-ltaS with PCR using primers YW58 and YW59, followed by Gibson assembly for gap closure. The removal of the *ltaS* coding region was confirmed with Sanger sequencing (Massachusetts General Hospital DNA Core). Plasmid pET-ltaS that mediates isopropyl β-D-1-thiogalactopyranoside (IPTG)-inducible over-expression of *ltaS* was generated through insertion of the *ltaS* coding region immediately after the IPTG-inducible promoter region of pET-28a(+) (Novagen^®^). Successful insertion was confirmed with Sanger sequencing (Massachusetts General Hospital DNA Core). Confirmed construct was transformed into *E. coli* BL21 (DE3) pLys for expression analysis.

### Antibiotic susceptibility testing

Antibiotic susceptibility testing was performed according to the BioMérieux E-test protocol with slight modifications. Specifically, a single colony of either the *S. mitis* ATCC 49456 wild type or *ΔltaS* strain was selected from cation-adjusted Mueller-Hinton (MH) (BD Bacto) agar cultures, inoculated into 1 mL of MH broth, and incubated for 6-8 hours at 37°C with 5% CO_2_. Then, 2 mL of fresh MH broth was added to the 1 mL culture, and the incubation was resumed. After overnight incubation, the OD_600nm_ of the cultures were measured, and samples having a value of OD_600nm_ < 0.2 were excluded from the following experimental procedures. Cultures were spread onto prewarmed MH agar plates with sterile cotton-tipped applicators, and plates were air dried for 15-20 minutes inside a biosafety cabinet. Then, E-test strips (ETEST® by BioMérieux) prewarmed to room temperature were applied to the plates with aseptic technique. The plates were incubated overnight at 37°C with 5% CO_2_. The minimum inhibition concentration (MIC) was determined by the intersection of the zone of inhibition with the E-test strip. At least three biological independent replicates were performed for each antibiotic-strain combination.

### Western blot analysis

Detection of Type I LTA via Western blot analysis was performed as previously described (39, 71).

For *E. coli*, single colonies of *E. coli* containing pET-ltaS were grown overnight in LB broth with 50 μg/ml kanamycin and 5 μg/ml chloramphenicol, followed by dilution to an OD_600nm_ of 0.1 with fresh media into two replicates. After 3 hours incubation at 37°C, IPTG was added to one set of cultures to a 1 mM final concentration, followed by another 30 minutes incubation at 37°C. Cell densities were normalized to an OD_600nm_ of 0.6, and 1 mL was pelleted, washed, resuspended in 100 μl 2× Laemmli sample buffer, and boiled for 15 min. Boiled samples were stored at −20°C prior to electrophoretic analysis.

For *S. aureus*, single colonies of each *S. aureus* strain were grown overnight in TS broth with 0.5 mM IPTG, 5 μg/ml erythromycin and 250 μg/ml kanamycin, and then sub-cultured to an OD_600nm_ of 0.1 into fresh TS broth containing 5 μg/ml erythromycin, 250 μg/ml kanamycin, and either 150 ng/ml ATC or 150 ng/ml ATC with 0.5 mM IPTG. After 3 hours incubation, the OD_600nm_ was measured, and cells equivalent to 1 ml of 1.2 OD_600nm_ were pelleted. Cell pellets were washed and resuspended with 1 ml phosphate buffered saline (PBS), followed by 5 cycles of bead-beating at 6.5 m/s for 45 seconds, with 5 minutes on ice between cycles (FastPrep-24™ MP Biomedicals). After centrifugation at 200 g for 1 min, cell lysates were collected, followed by pelleting at 17000 g for 10 minutes. The material was resuspended in 100 μl 2× Laemmli sample buffer (Bio-Rad) followed by boiling for 15 minutes in a heating block.

For streptococci and *E. faecalis*, unless indicated, OD_600nm_ values of the overnight cultures were measured, followed by pelleting of cells equivalent to 1ml of 1.2 OD_600nm_. Induction of *ltaS* overexpression in *S. mitis* was conducted similarly as in *S. aureus*. Specifically, overnight cultures of *S. mitis* containing either pitetR-ltaS or pitetR were diluted to an OD_600nm_ value of 0.1 into fresh TH broth with 150 ng/ml ATC. After 7 hours incubation, cells equivalent to 1 ml of 1.2 OD_600nm_ were harvested. All cell pellets were washed and resuspended with 1 ml PBS, then followed with the same cell disruption and lysate preparation processes as described above for *S. aureus* samples.

Separation of cell lysate materials are conducted through sodium dodecyl sulfate - polyacrylamide gel electrophoresis (SDS-PAGE). Specifically, 15 μl of each boiled sample was loaded to a 15% SDS-PAGE gel, followed by electrophoresis at consistent 100 voltage and subsequent PVDF membrane transfer at consistent 350 mA. The blocking solution was PBS containing 0.05% (w/v) Tween 20 and 10% (w/v) non-fat milk; antibody solutions were PBS with 0.05% (w/v) Tween 20 and 5% (w/v) non-fat milk. For *S. aureus* samples, 3 μg/ml human IgG (Sigma) was added to the blocking and antibody solutions to block the activity of protein A. Primary antibody targeting Type I LTA (clone 55, Hycult Technology) and secondary antibody (HRP-conjugated anti-mouse IgG, Cell Signaling) were used at dilutions of 1:2500 and 1:5000 respectively. After adding HRP substrate (Immobilon^®^ Western, Millipore) and shaking at room temperature for 3 minutes, chemiluminescence signals were detected with the ChemiDoc™ Touch Imaging System (Bio-Rad) with default Chemiluminescent settings. Relative band intensity was analyzed with the Image Lab Software (Bio-Rad).

### Lipidomics analysis

Extraction of total lipids from stationary phase cells was performed by acidic Bligh-Dyer extraction as previously described (18). Specifically, cells were grown to stationary phase, followed by collection and storage at −80°C until lipid extraction with the acidic Bligh-Dyer methods. The dried lipid extracts were dissolved in a mixture of chloroform and methanol (2:1, v/v) before LC/MS analysis. NPLC-ESI/MS of lipids was performed as previously described (41, 72) using an Agilent 1200 Quaternary LC system (Santa Clara, CA) coupled to a high resolution TripleTOF5600 mass spectrometer (Sciex, Framingham, MA). An Ascentis^®^ Si HPLC column (5 μm, 25 cm × 2.1 mm, Sigma-Aldrich) was used. Mobile phase A consisted of chloroform/methanol/aqueous ammonium hydroxide (800:195:5, v/v/v). Mobile phase B consisted of chloroform/methanol/water/ aqueous ammonium hydroxide (600:340:50:5, v/v/v/v.). Mobile phase C consisted of chloroform/methanol/water/aqueous ammonium hydroxide (450:450:95:5, v/v/v/v). The elution program was as follows: 100% mobile phase A was held isocratically for 2 min and then linearly increased to 100% mobile phase B for 14 min and held at 100% B for 11 min. The LC gradient was then changed to 100% mobile phase C for 3 min and held at 100% C for 3 min, and finally returned to 100% A over 0.5 min and held at 100% A for 5 min. Instrumental settings for negative ion ESI and MS/MS analysis of lipid species were as follows: ion spray voltage (IS) = −4500 V; current gas (CUR) = 20 psi (pressure); gas-1 (GS1) = 20 psi; de-clustering potential (DP) = −55 V; and focusing potential (FP) = −150 V. The MS/MS analysis used nitrogen as the collision gas. Data acquisition and analysis were performed using the Analyst TF1.5 software (Sciex, Framingham, MA).

### Serum survival test

Overnight bacterial cultures were pelleted and washed with PBS, followed by sub-culturing into different media to an OD_600nm_ of 0.1. Cultures were incubated at 37°C with 5% CO_2_ for 8 hours. At t=0 and t=8 hours of incubation, viable bacterial cells were determined by serial dilution and plating on TH agar plate.

## Acknowledgements

We gratefully acknowledge Dr. Angelika Gründing for providing the *S. aureus* strain ANG113 and ANG499.

This work was supported by grant R21AI130666 from the National Institutes of Health and the Cecil H. and Ida Green Chair in Systems Biology Science to K.P, grant R56AI139105 and R01AI148366 from the National Institutes of Health to K.P and Z.G., and U54GM069338 to Z.G.

---

**Fig. S1.**
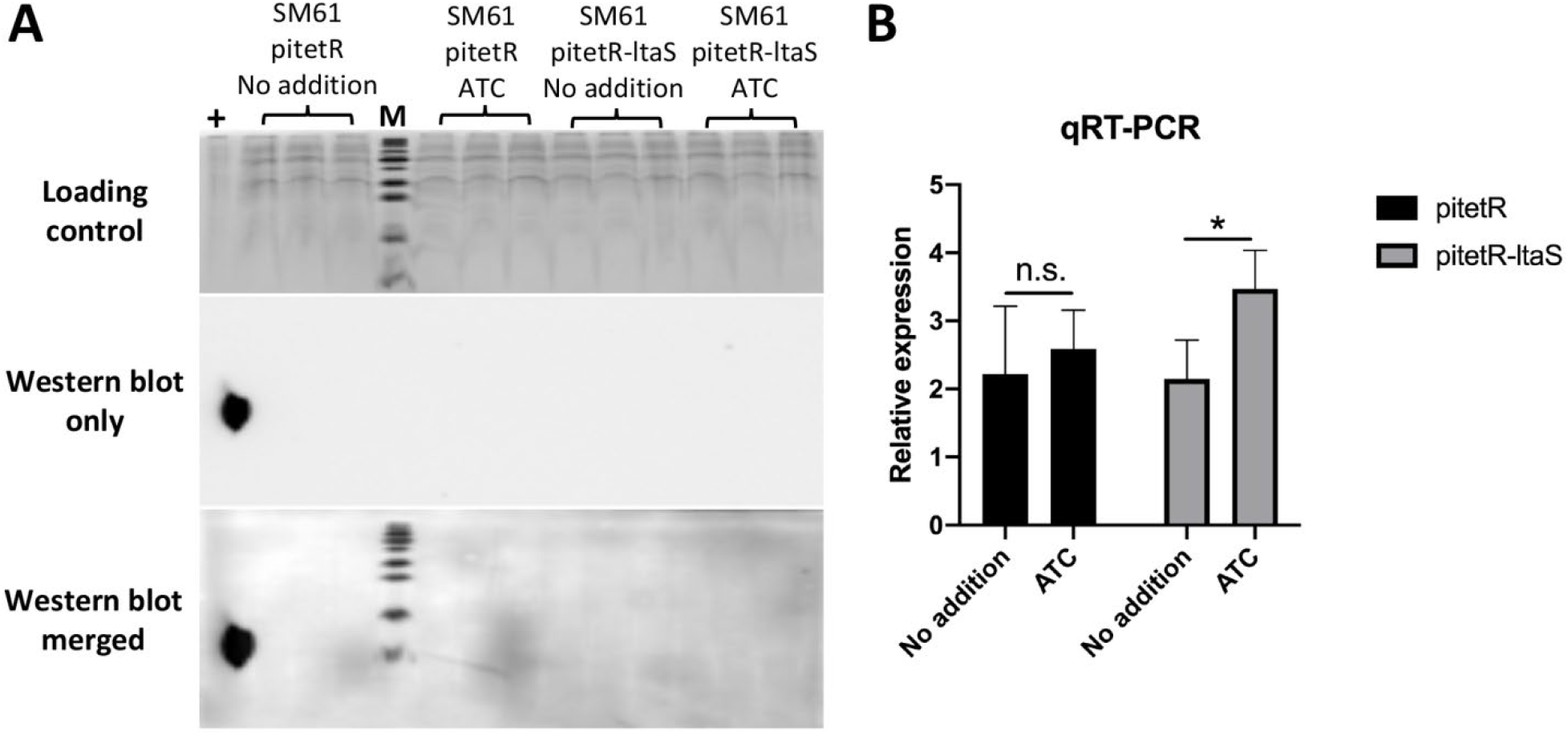
A) Western blot detection of Type I LTA in *Streptococcus mitis* ATCC 49465 (SM61) containing either the *ltaS* expression plasmid pitetR-ltaS or empty plasmid control (pitetR). Over-expression of *ltaS* was induced by addition of 150 ng/ml anhydrotetracycline (ATC). Cell lysates were prepared from cultures grown to stationary phase. Cell lysate of *S. aureus* was used as positive control (+). Western blot figure was obtained 2 minutes after the saturation of the positive control signal. Loading control was stained with Commassie blue. B) qRT-PCR detection of the transcript levels of *ltaS* from SM61 containing pitetR-ltaS or pitetR with or without ATC induction. Total RNA was harvested from mid-log phase cells exposed to ATC for 30 minutes. Relative expression levels of *ltaS* were normalized to that of 16S rRNA of the same sample with the ΔΔCt method. Four biologically independent replicates were obtained for each sample. Statistical analysis was performed with one-way ANOVA; “*” indicates 0.01 < *P*-value < 0.05 and “n.s.” indicates non-significant.

**Fig S2.**
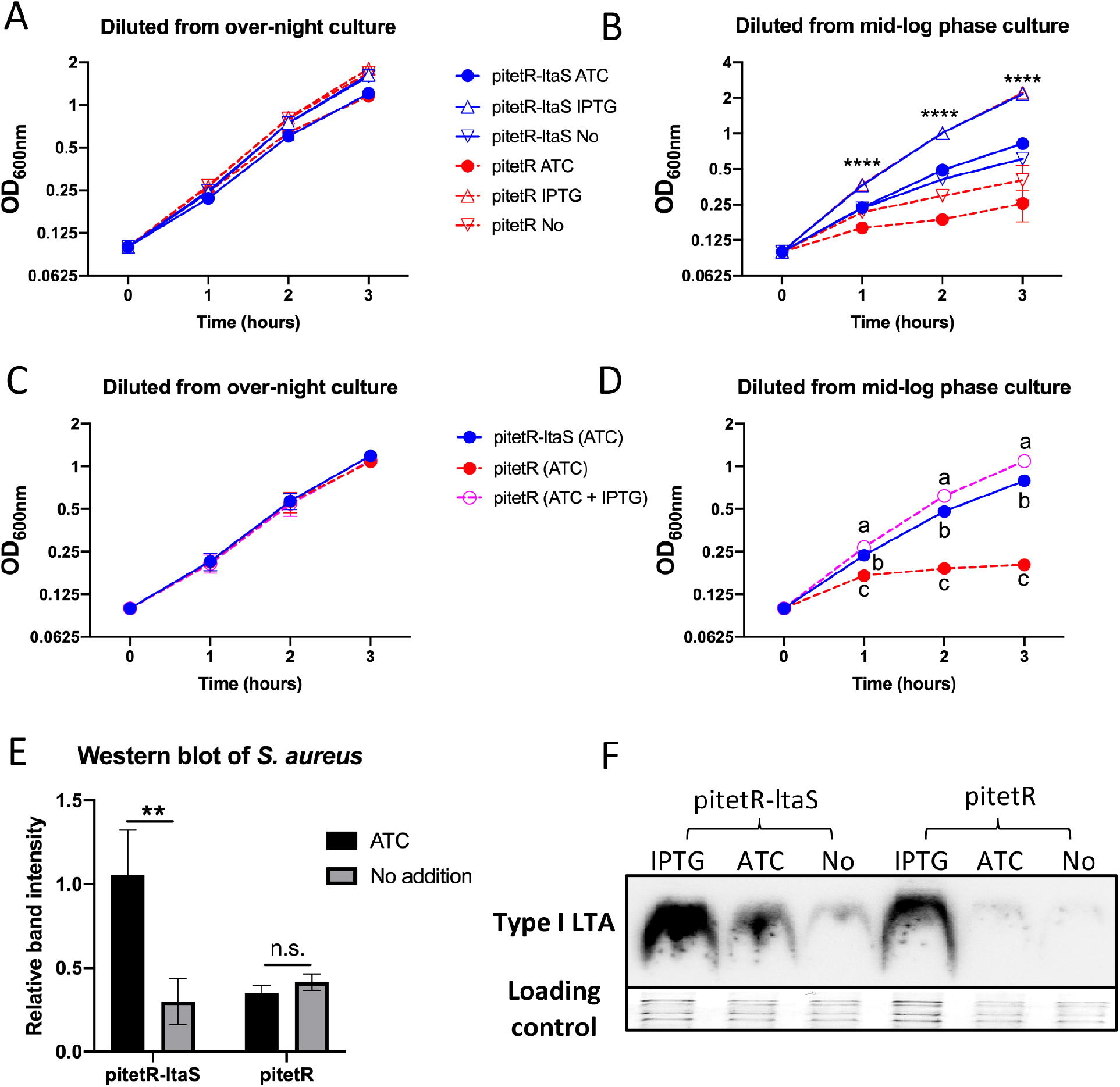
*S. mitis ltaS* complements the function of *S. aureus ltaS*. Single colonies of *S. aureus* ANG499 strain containing either pitetR-ltaS or pitetR were grown over-night in Trypic Soy broth (TSB) followed by dilution into 0.1 of OD_600nm_ with fresh TSB with indicated addition of 150 ng/ml ATC, 0.5mM IPTG, no addition (No), or both 150 ng/ml ATC and 0.5mM IPTG (ATC + IPTG). Values of OD_600nm_ were measured every hour for the first 3-hour incubation (A & C). Then, cells equal to 1ml of 1.2 OD_600nm_ were harvested from each sample for Western blot analysis (E & F); in the same time, another dilution same as describe above was performed, followed with continued incubation for another 3 hours and measurement of OD_600nm_ values every hour (B & D). For E, band intensities were normalized to that of sample pitetR-ltaS (ATC) and loading control. Data shown as plots were obtained from four biologically independent replicates. Statistical analyses were performed with one-way ANOVA. Significant differences were shown as “**” for 10^−1^ < *P*-value < 10^−2^, “****” for *P*-value < 10^−6^. Non-significant is indicated by “n.s.”. For panel D, at time point 1, 2, and 3, statistically different groups were indicated separately as “a”, “b”, and “c”. *P*-values between each group are all < 10^−6^.

**Fig. S3.**
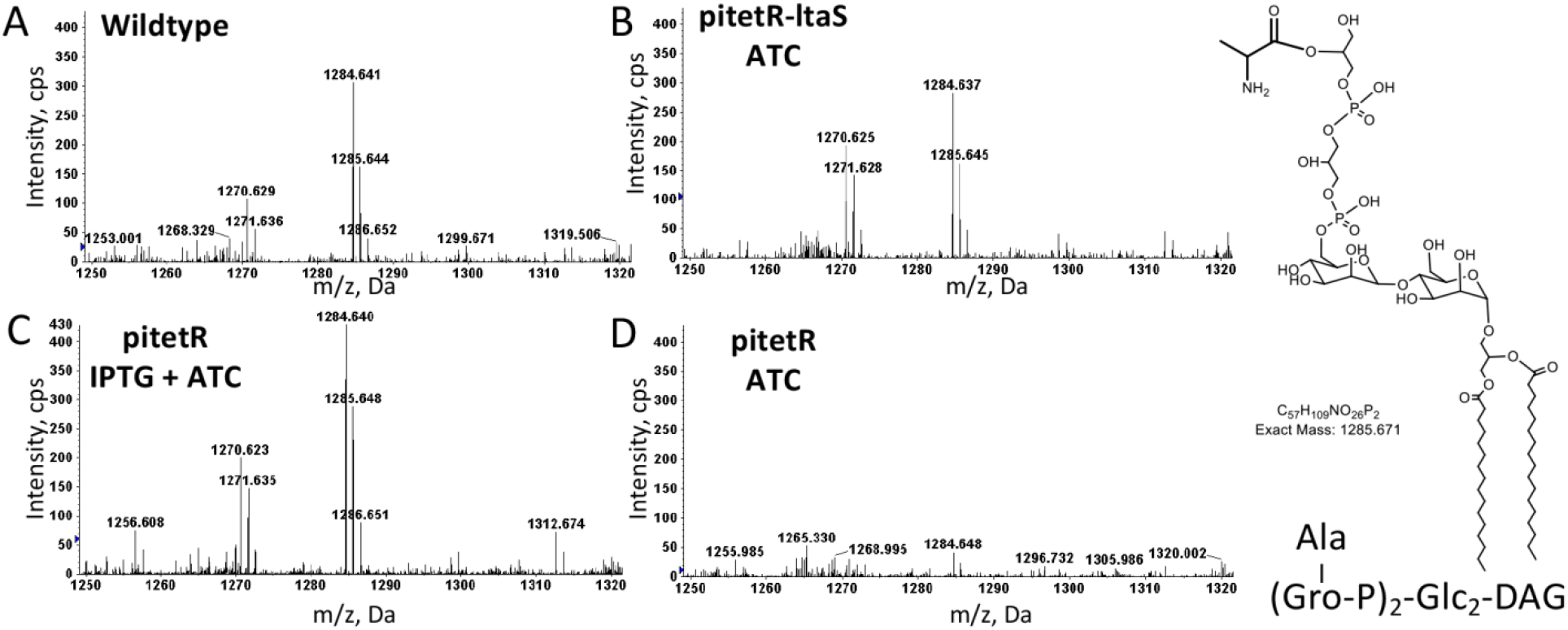
MS detection of Type I LTA intermediate that contains two Gro-P units with one Ala modification in the lipid extracts of *S. aureus*. *S. aureus* wildtype (A) and native *ltaS* deficient strain containing either plasmid pitetR-ltaS (pitetR-ltaS) (B), or empty vector control (pitetR) (C & D) were incubated to stationary phase in Tryptic Soy broth with the addition of either 150 ng/ml anhydrotetracycline (ATC) or 150 ng/ml ATC with 0.5 mM IPTG as indicated. Total lipids were extracted with a modified acidic Bligh-Dyer extraction method and analyzed with NPLC-ESI/MS in the negative ion mode. Shown are the deprotonated [M-H]^−^ ions of Type I LTA intermediate containing two Gro-P units with one Ala modification (retention time: ~23.5-24.0 min; most abundant *m/z* 1284.6). Abbreviations: Ala, alanine; Gro-P, glycerophosphate; Glc, glucosyl; DAG, diacylglycerol.

**Fig. S4.**
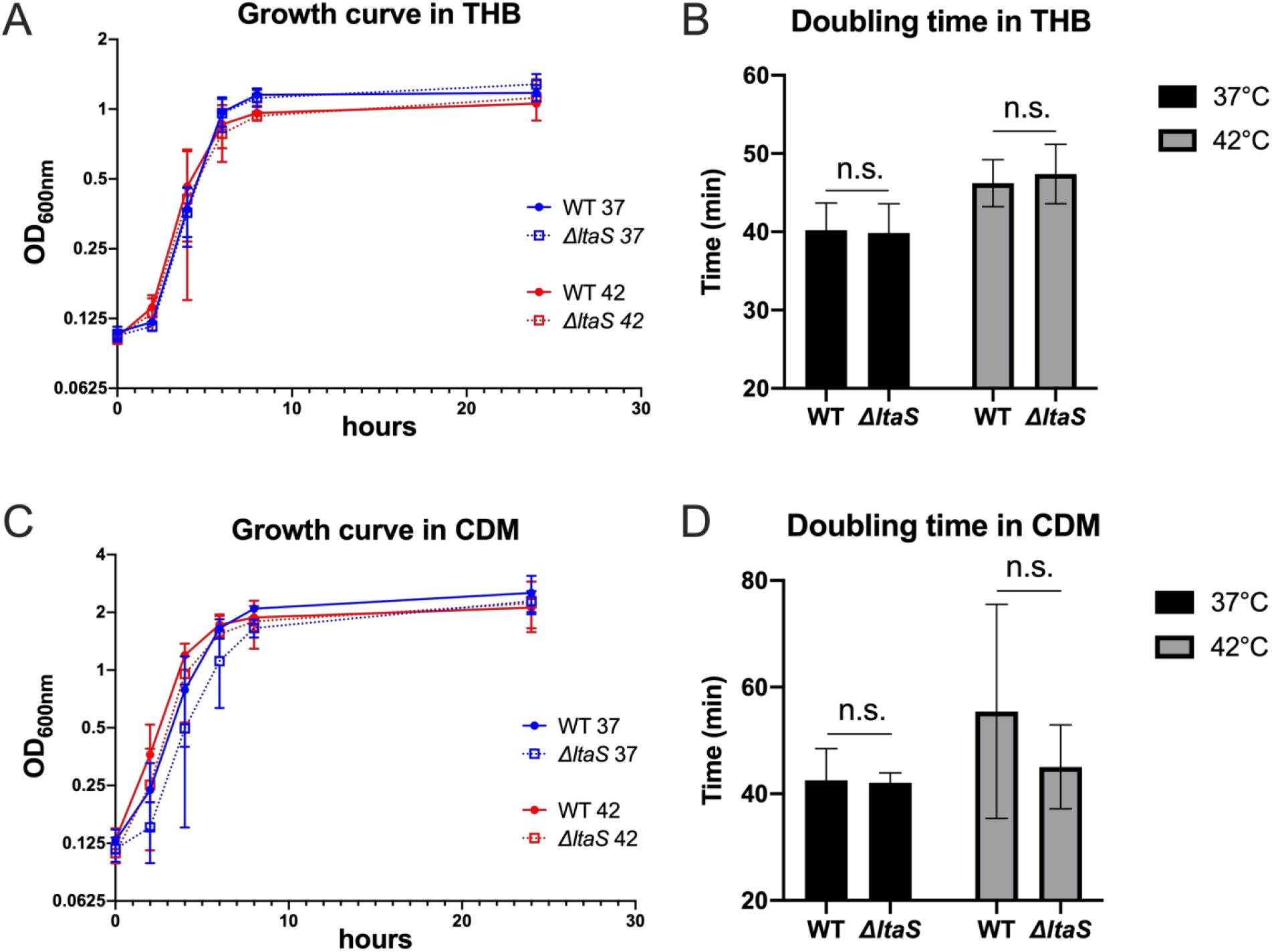
Growth curve of *S. mitis* under different culture conditions. Single colony of either *S. mitis* ATCC 49456 (SM61) wildtype (WT) or *ΔltaS* strain cultured overnight in either Todd Hewitt broth (THB) or chemically defined medium (CDM) were diluted into fresh indicated medium to an OD_600nm_ value of 0.1, followed by incubation at either 37°C (blue lines) or 42°C (red lines) as indicated in A & C. Values of OD_600nm_ were measured at incubation time of 0, 2, 4, 6, 8, and 24 hours. Doubling time shown in B & D was calculated using the OD_600nm_ values acquired at incubation time of 2, 4, and 6 hours. Data presented are mean values from either at least four (THB) or two (CDM) biological replicates, with standard deviations represented by the error bars. Statistical analyses were performed with One-way ANOVA; significant difference was determined by *P*-value < 0.05; “n.s.” represents statistical non-significant.

**Table S1.**
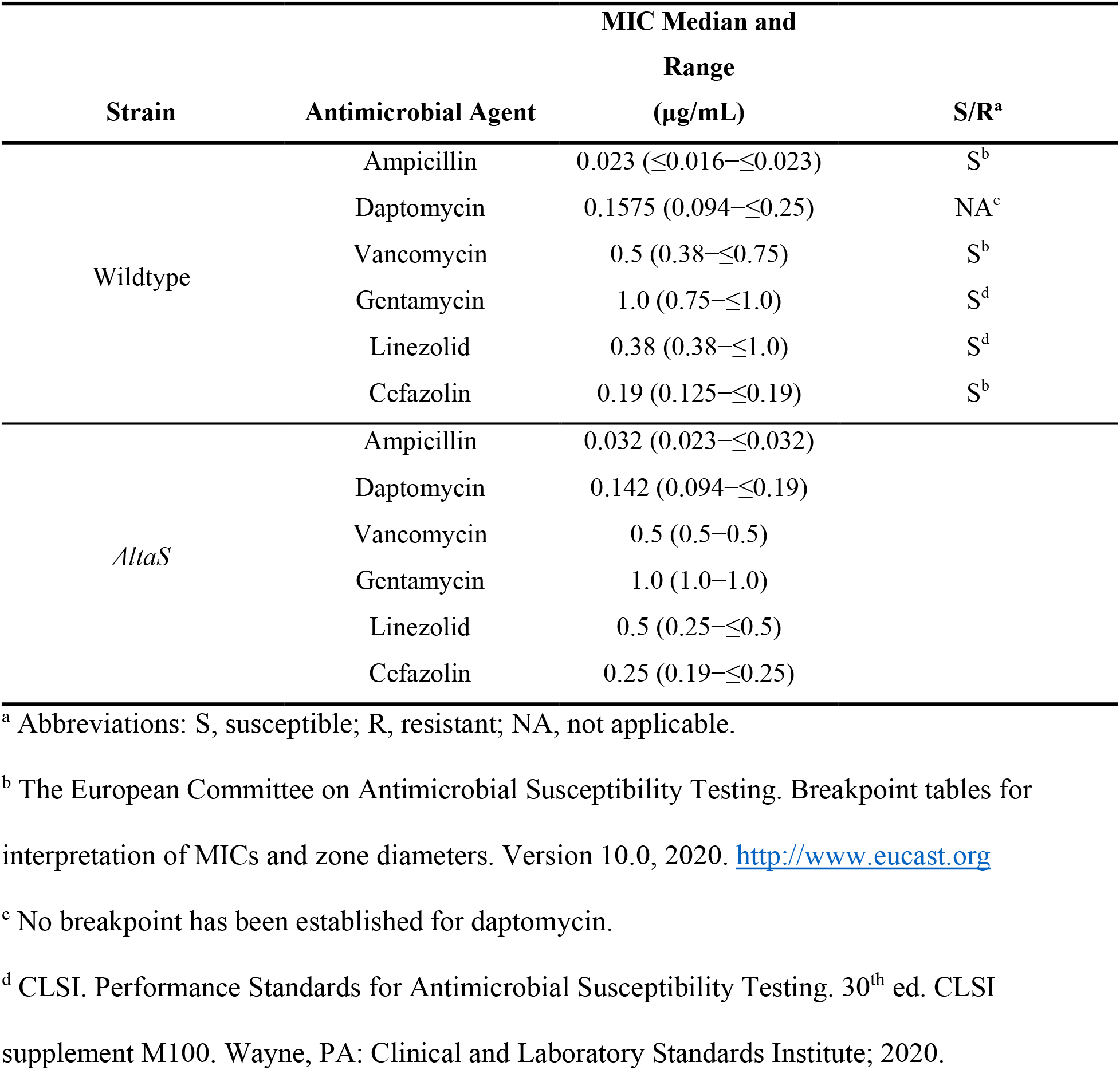
E-test results of the wildtype and *ΔltaS* strain of *S. mitis* ATCC 49456 (SM61)

**Table S2.**
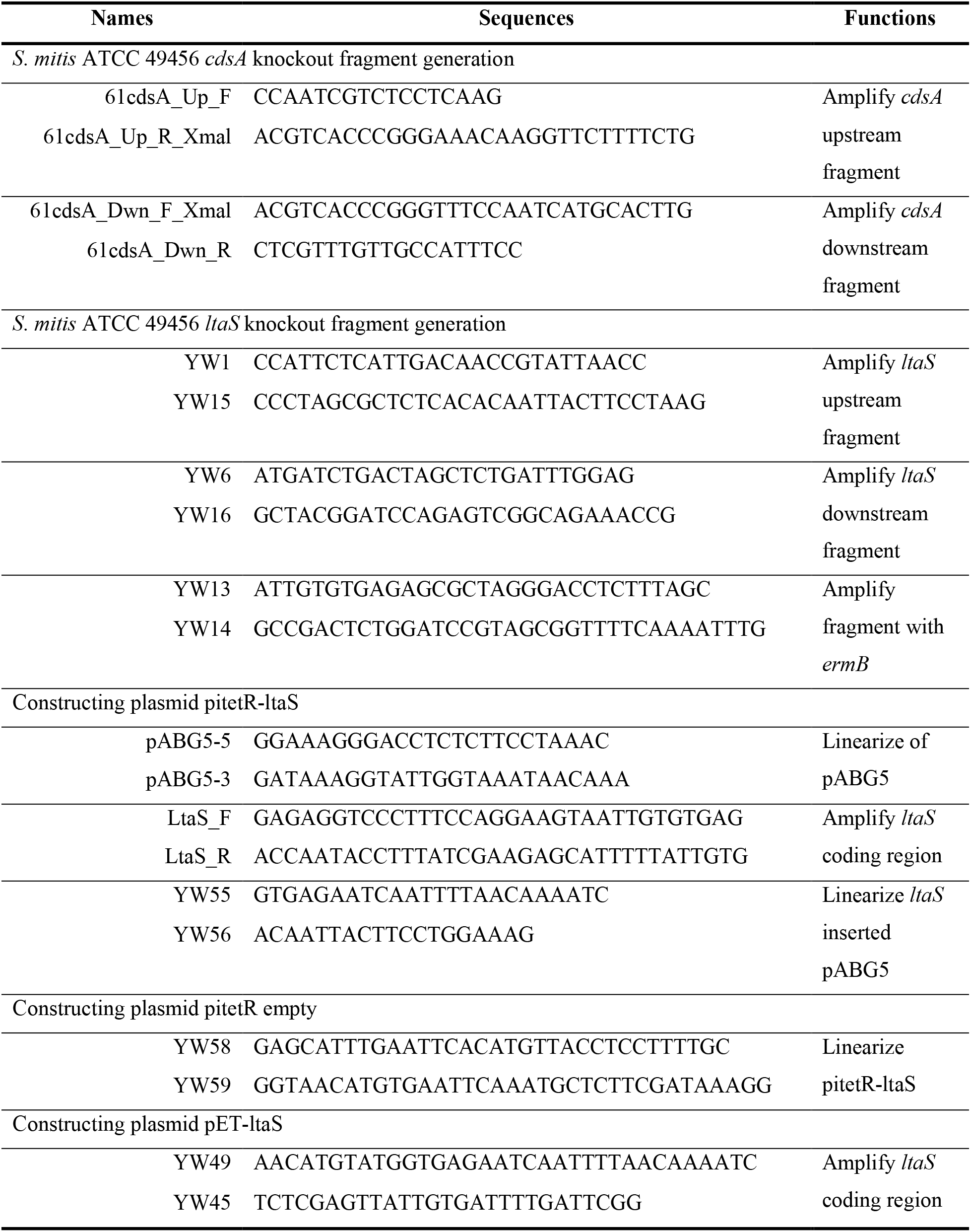
Primers used in this research

## Notes

### Competing Interest Statement

The authors have declared no competing interest.

